# CALISTO: A Standardized and Open-Source Software for Magnetic Tweezer Force Calibration

**DOI:** 10.64898/2026.10.17.759816

**Authors:** Stefanie D. Pritzl, Alptuğ Ulugöl, Caroline Körösy, Leonhard R. Schatt, Marek Vlasak, Koen R. Storm, Laura Filion, Jan Lipfert

## Abstract

Magnetic tweezers (MT) enable multiplexed single-molecule force spectroscopy while applying controlled forces and torques to biomolecules. They resolve forces below 1 pN and maintain constant forces over extended periods, but accurate calibration remains challenging due to drift, tracking artifacts, camera noise, and bead variability.

Here, we present CALISTO (CALIbration Suite for magnetic Tweezer Operation), an open-source software for MT force calibration. CALISTO analyzes position traces of DNA-tethered magnetic beads and surface-bound reference beads for drift correction. It implements three thermal-motion-based calibration methods: power spectral density, Allan variance, and Hadamard variance, with the latter providing increased robustness against drift and correlated measurement noise.

The graphical user interface combines automated analysis with user-guided quality control, including reference bead selection, z-offset determination, magnet-position classification, bead-wise force estimation, and tether extension analysis. Together, these steps generate a master force calibration curve. We systematically evaluated their effects on calibration performance.

The largest improvement resulted from automated filtering of problematic force plateaus using time-series characterization to identify traces affected by drift, tracking artifacts, or irregular fluctuations. Filtering reduced scatter and improved force estimation while preserving the underlying force–magnet position relationship. CALISTO thus provides a standardized, reproducible framework for reliable magnetic-tweezers force calibration.

**STATEMENT OF SIGNIFICANE:** Magnetic tweezers enable highly sensitive and long-duration measurements of forces and torques in single biomolecules, but their accuracy critically depends on reliable force calibration. CALISTO provides an open-source, standardized framework that integrates force calibration with automated quality control and quantitative assessment of experimental data. By identifying problematic measurements caused by drift, tracking artifacts, or irregular bead fluctuations, CALISTO substantially improves calibration reliability while preserving the underlying force–position relationship. The software implements complementary thermal-motion-based calibration methods and combines automated processing with user-guided validation in an accessible graphical interface. CALISTO facilitates reproducible magnetic-tweezers experiments across different experimental conditions and provides a practical foundation for more reliable quantitative single-molecule force measurements.

## INTRODUCTION

Single-molecule force spectroscopy (SMFS) techniques have emerged as powerful tools for probing the mechanical behavior of biological macromolecules under applied forces and torques, offering insights into diverse processes such as molecular motor activity, DNA and protein mechanics, and receptor-ligand interactions.^1-18^ Among the various SMFS platforms including optical tweezers and atomic force microscopy, magnetic tweezers (MT) offer distinct advantages.^1, 5, 6, 19-21^ MT enable the application of stable forces over a broad range (∼0.01–100 pN) and across timescales from milliseconds to days ^22, 23^, while allowing for high-throughput, multiplexed measurements of hundreds of tethered molecules under uniform force conditions.^22, 24-27^ In particular, MT allow access to forces in the sub-piconewton range (≤1 pN), which is essential for detecting conformational changes and weak molecular interactions relevant to cellular functions such as motility, differentiation, and development.^28-31^ They achieve this without requiring feedback control^12^ and without introducing heat or photodamage^32^. Furthermore, specific magnet geometries allow for precise application and measurements of torque at the single-molecule level.^33-35^

In MT, molecules of interest are tethered between a functionalized surface and micrometer-sized superparamagnetic beads. External magnets apply forces and torques to control the extension and twist of the bead’s tether, enabling manipulation and observation of the molecular system (Figure 1A). State-of-the-art cameras enable imaging large fields of view (≥ 0.5 mm^2^), allowing for simultaneous tracking of tens to hundreds of beads and facilitating multiplexed measurements under uniform conditions (Figure 1B). A key requirement in these experiments is accurate force calibration, typically based on analyzing the thermal fluctuations of tethered beads using the equipartition theorem (Figure 1C).^36-38^ According to this method, the magnetic bead-tether system can be modelled as a harmonically trapped particle with trap stiffness *k*_*i*_ along axis *i*∈{*x,y,z*}that is related to the real-space variance (RSV) *σ*^2^ of the bead’s thermal fluctuations via:

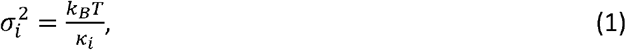

where *k*_*B*_ is the Boltzmann constant and *T* is the absolute temperature. One can show that the trap stiffness in the transverse direction (in which the magnetic field constrains the rotation of the bead) is given by *K*_‖_=*F*_*Z*_/⟨*L*_*ext*_⟩, so that the axial force F is given by^38^:

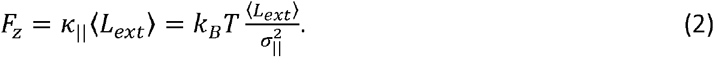

with ⟨*L*_*ext*_⟩representing the mean extension of the tether and 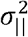 the variance of the transverse fluctuations in the direction parallel to the magnetic field. For accurate force calibration, the challenge is, therefore, to determine the mean extension of the tethers and the variance of the transverse fluctuations accurately and without bias.

**Figure 1:**
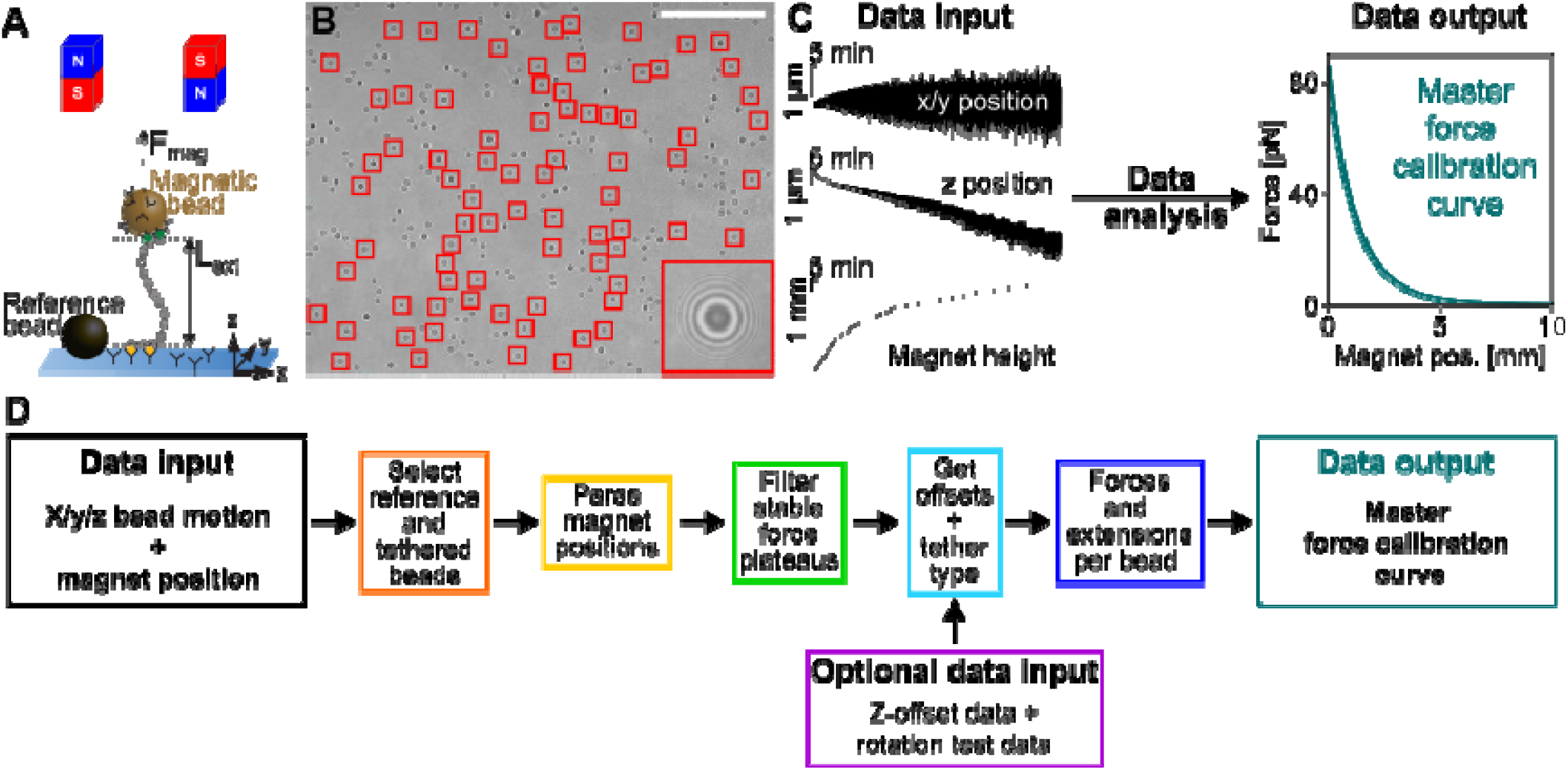
Overview of the magnetic tweezers setup and calibration principle. **A)** Schematic of a MT setup. A DNA molecule with extension ***L***_*ext*_ is tethered between a functionalized surface and a superparamagnetic bead. External magnets apply a magnetic force ***F***_*mag*_ to the bead. Surface-attached reference beads are tracked for drift correction. B) Representative field of view from the MT setup, with red squares marking 78 beads tracked over time. The inset shows a magnified view of a bead’s diffraction pattern used for position tracking. Scale bar: 100 µm. Adapted from ref. ^51^ C) In CALISTO, position fluctuation traces of DNA-tethered beads in the x- or y-, and z-directions and the corresponding magnet heights are analyzed to generate the final force calibration curve, i.e. force vs. magnet position. D) Flowchart of the calibration pipeline. CALISTO takes bead position traces and corresponding magnet-position data as input, with optional z-offset and rotation test files. The workflow includes bead identification and classification, magnet-position parsing, detection of stable force plateaus, z-offset determination, and screening for tether quality. From these steps, force–magnet position and force– extension relationships are derived for individual beads and combined to generate a global force calibration curve.

Long double-stranded DNA (dsDNA) molecules with contour lengths exceeding 1 µm are commonly used for force calibration due to their well-characterized mechanical properties. One advantage of using DNA tethers is that their force-extension response is well known: at low forces, their force-extension behavior is accurately described by the worm-like chain model^39-41^ and they exhibit a distinct overstretching transition at ∼65 pN^42-44^. Using long (≥*µm* ) tethers facilitates precise measurement of the bead’s thermal motion compared to shorter tethers: tracking errors are less significant, fluctuations are slower reducing artifacts from camera exposure time, and off-center tethering effects are minimized, compared to short tethers.^25, 38, 45-50^

While, in principle, forces can directly be calibrated from measurements of the mean extension and the variance of the transverse fluctuations via Eq. (2), in practice the RSV-based approach should be avoided, as it is sensitive to common experimental artifacts. Downsampling, due to finite camera exposure times can blur trajectories and artificially reduce the variance, in particular at high forces where the characteristic time scales of the fluctuations are short.^48^ Additional noise from electronic components, optical artifacts, or tracking errors can introduce artifactual fluctuations;^51^ in particular residual mechanical drift, even after reference bead correction, remains a challenge in MT, which all tend to artificially increase the apparent variance.

To overcome these limitations and to provide a standardized calibration workflow, we developed CALISTO, a user-friendly, open-source Python-based graphical user interface (GUI) for force calibration in multiplexed MT experiments. CALISTO integrates thermal motion-based calibration methods, including power spectral density (PSD)^52^, Allan variance (AV)^52^, and Hadamard variance (HV)^51^. Of these, HV is the most robust against strongly correlated noise and drift by using a second-order finite difference approach that cancels linear drift contributions and enhances frequency resolution.^51^ Simulations and experimental data have previously demonstrated that HV improves calibration accuracy under realistic conditions with residual drift and low signal-to-noise ratios.

The software guides users through each step of the calibration workflow, from importing raw data to the generation of a final force vs. magnet position calibration curve (Figure 1C). A flowchart of the full workflow is shown in Figure 1D: CALISTO requires bead position traces and corresponding magnet-position data as input, with optional z-offset and rotation test files to define the absolute z-axis origin (typically setting the z-position where the bead touches the flow cell surface to zero) and assess tether integrity, respectively. Data analysis includes bead identification and classification (reference or tethered), parsing of magnet-position data, detection of stable force plateaus (i.e. of the segments of the time trace that correspond to a given magnet position and thus force), z-offset determination, and screening for multiple tethers or off-center attachments. From these steps, force–magnet position and force–extension relationships are obtained for individual tethered beads and combined to construct a global force calibration curve using only well-characterized beads.

By standardizing the calibration process, CALISTO reduces user-induced variability and enhances the reproducibility of MT-based SMFS experiments. Its modular design and open-source implementation facilitate adaptation to different experimental setups. CALISTO therefore provides a robust and consistent framework for force calibration in magnetic tweezers.

## MATERIALS AND METHODS

To develop CALISTO, a standardized, open-source graphical user interface (GUI) for magnetic tweezer force calibration, we combined analytical and numerical approaches and validated the performance of the GUI implementation against experimental data. In the following sections, we outline the required data inputs, describe the computational methods implemented in the GUI, and present the magnetic tweezer setup and calibration protocol. CALISTO is implemented in Python 3.12 and is available freely at github.com/lipfertlab/calisto.

### Data Requirements for CALISTO Analysis

CALISTO is used to analyze experimental data from MT force calibration measurements. The software requires a set of structured input data that contain the bead motion and the movement and position of the external magnets. The bead motion input consists of position time traces, where bead coordinates in the x, y, and z directions are recorded together with a time column. The motor data input must specify the magnet position (height) – typically relative to the flow cell surface – for each time point or step, to link the bead response to the applied magnetic field. These data inputs form the basis for identifying force plateaus and extracting the force–magnet position relation. The data can be provided either as plain text or HDF5 files. For plain text input, bead position traces and motor-position data must be supplied as separate files. In contrast, for HDF5 input, both datasets must be stored within the same file following the required internal structure. Detailed specifications of the file formats and organization are provided in the CALISTO GitHub repository (github.com/lipfertlab/calisto).

An optional but recommended input is a short measurement to determine the z-offset (“z-offset test”), and thus the bead–surface coordinate system and the origin of the z-axis. Although z-positions are measured on an absolute scale via a look-up-table, the origin remains a convention. CALISTO adopts the convention to set the bead’s z-position when it contacts the surface as *z* = 0. The z-offset is experimentally determined from a continuous trace per bead in which the force is first set to a high value (≥10 pN), then reduced to near zero (∼0 pN), and subsequently increased again to a high value (≥10 pN). Accurate z-offset determination is essential for obtaining physically meaningful DNA extensions. Omitting this step leads to incorrect force estimates and can result in unphysical values, such as negative extensions.

Finally, a rotation test is recommended to assess tether integrity and ensure that only beads tethered by a single double-stranded DNA molecule are included in the analysis. This corresponding file must contain tether extension measurements under controlled magnet rotations, with the corresponding magnet position and rotation information provided in the motor file. The detailed interpretation of these traces is described below in the user manual. Briefly, at intermediate force (∼3-5 pN), negative rotations (−70 turns), which underwind the DNA helix, are applied: single DNA tethers show no change in extension due to torque-induced melting, whereas multiple tethers form braids and shorten and can be excluded from further analysis. At low force (∼0.4 pN), positive rotations (+70 turns) overwinding the DNA helix are applied to detect whether the DNA molecule is supercoilable. Intact, torsionally constrained tethers form plectonemic supercoils leading to a reduction of their extension, while nicked tethers show no change in tether extension. Together, these inputs ensure that calibration is performed on physically consistent and well-defined datasets.

### Automated Selection of Good Reference Beads for Global Mechanical Drift Correction

Reliable drift correction requires tracking multiple reference beads that are firmly attached to the surface and not tethered to a DNA. In practice, we recommend using at least three reference beads and, where possible, approximately five to ten reference beads to ensure robust drift correction, consistent with previous analyses of multiplexed magnetic-tweezers measurements.^53^ These beads should be recorded under the same acquisition conditions as the tethered beads and over the full duration of the experiment to capture and correct for mechanical drift. The full position traces of reference and tethered bead traces must be stored in the same file (see GitHub repository for detailed information on file structures: github.com/lipfertlab/calisto).

To automate the selection of suitable and artifact-free reference beads for global mechanical drift correction from the pool of initially identified candidates (user input in the software, Figure 2), which we refer to as high-quality reference beads, we implement a four-step procedure comprising (i) data smoothing, (ii) outlier detection, (iii) mean trace extraction and detrending, and (iv) final reference bead selection. Before the final selection, we perform additional iterative outlier detection steps to refine the dataset, ensuring that the selected traces are free from artifacts and systematic deviations. The details of each step are described below.

**Figure 2:**
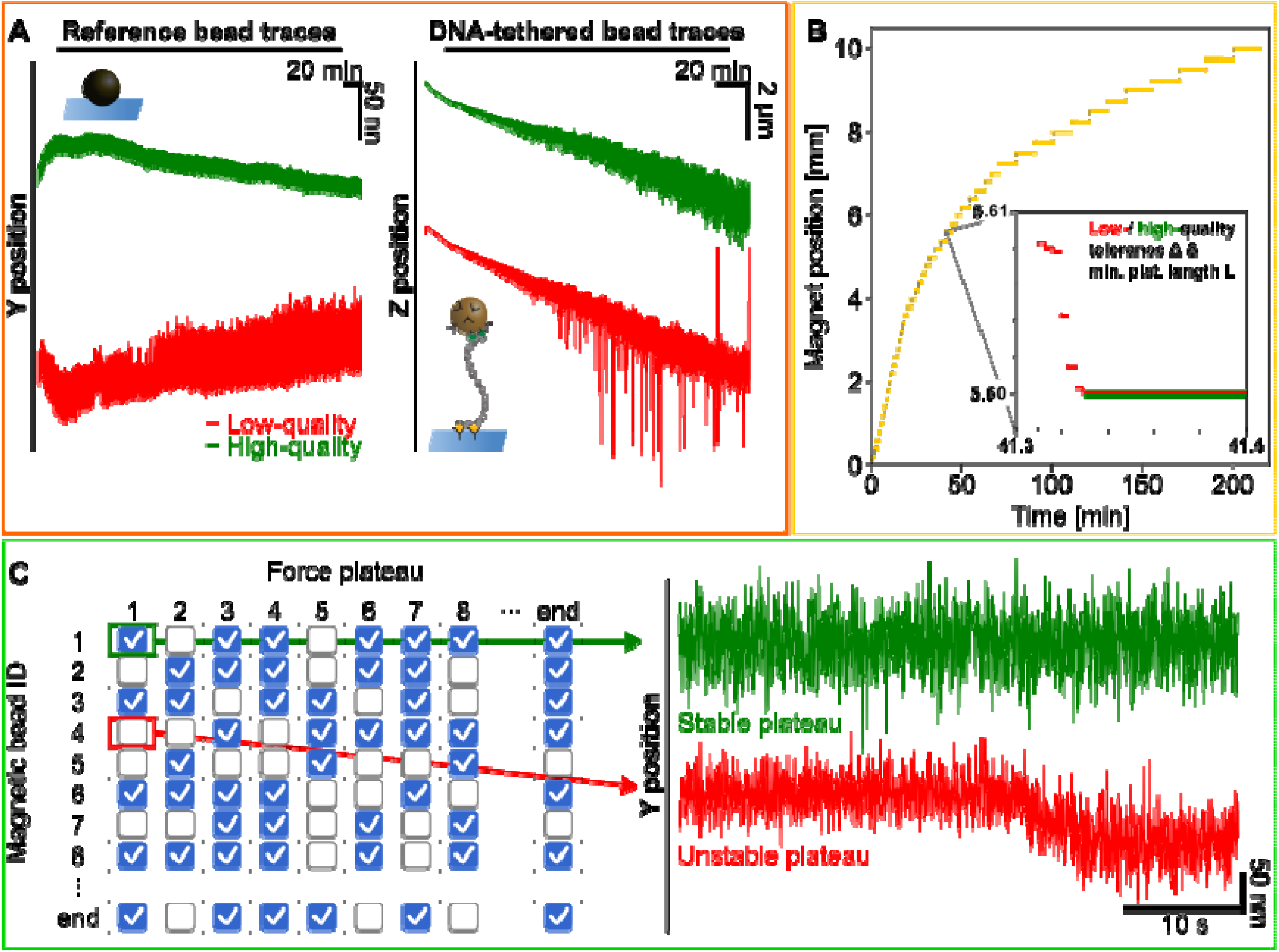
Bead selection, magnet-position classification, and plateau detection for force calibration. **(A)** Example traces of reference and DNA-tethered beads. A low-quality reference bead (red) shows a higher noise level compared to a high-quality one. Likewise, a low-quality tethered bead trace (red) deviates from the mean and exhibits elevated spikes in the z-direction, unlike the stable trace (green). (B) Time-lapse motor movement showing 49 magnet positions. Inset: Comparison of effective (green) and poor (red) settings for tolerance threshold and minimum plateau length. Correct settings detect only the true stationary position at 5.6 mm, while poor settings include spurious parts of the trace caused by e.g. magnet movement. (C) Example of how stable segments or plateaus in time traces are selected. Left: checkmarks indicate stable force plateaus; empty entries mark unstable ones. Left: representative stable (green) and unstable (red) force plateau position time traces.

#### (i) Smoothing of Reference Traces

To process the input reference bead traces, we apply an exponential smoothing method to the differences between successive data points along the x-, y-, and z-axes independently. The smoothing is automated through the Simple Exponential Smoothing (SES) algorithm^54^. This approach helps to reduce high-frequency fluctuations while preserving essential features of the reference traces.

#### (ii) Outlier Detection

To identify the reference traces that deviate from a common trend, we perform an outlier detection using the isolation forest algorithm, a robust unsupervised learning method for anomaly detection.^55, 56^ This algorithm is applied to the smoothed reference traces to identify and mask outliers. The reference beads that are marked as outliers are excluded in the remaining steps.

#### (iii) Mean Trace Extraction and Detrending

To obtain a representative mean drift trend, we compute the average over all non-outlier reference traces along each axis (x, y and z). Exponential smoothing is then applied independently to each x-, y-, and z-position axes to minimize fluctuations. The smoothed mean trace is subsequently subtracted from individual reference traces to remove slow variations and drift, yielding a first set of detrended traces. After detrending, well-behaved traces should fluctuate around zero without systematic drift.

#### (iv) Final Selection of Reference Beads

To further refine the reference trace selection, we repeat step (ii) using the reference traces picked in step (iii). Then, we calculate the mean reference trace by averaging the raw, not-smoothed, traces of the non-outlier reference beads. This averaged trace is finally used to correct the tethered bead traces for drift.

### Noise Classification for DNA-Tethered Bead Trace Selection

To obtain reliable force estimates, it is essential that each force plateau corresponds to a stationary trace whose fluctuations are dominated by the thermal motion of the bead. In practice, some plateaus contain additional drift, tracking artifacts, or disturbances caused, for example, by imperfect bead tracking, mechanical instabilities in the MT setup, or changes in focus. These effects introduce additional (correlated) fluctuations and trends into the trace and can strongly bias variance-based estimates of the magnetic trap stiffness. We therefore identify and exclude plateaus whose fluctuation properties are inconsistent with the thermal motion of a trapped bead.

For each DNA-tethered bead trace, the stable force plateaus corresponding to constant magnet positions are analyzed independently. The dominant noise behavior is characterized using a power-law noise model, in which the power spectral density scales as *S*(*f*) *∝* 1/*f*^*α*^ . Different values of the spectral exponent *α* correspond to different types of stochastic fluctuations, e.g. *α*=0 corresponds to Gaussian fluctuations with no temporal correlations, and *α*=2 to free diffusion. Values of *α* > 2 indicate increasingly strong low-frequency variations and are associated with nonstationary, drift-like behavior. In contrast, negative values of *α* indicate that the fluctuations become stronger at higher frequencies, which is consistent with high-frequency-dominated or anti-correlated fluctuations and may arise from localization noise, instrumental noise, or data-processing artifacts.

To estimate *α*, we use the lag-1 autocorrelation method that measures the correlation between consecutive datapoints in the trace.^57^ For power-law noise processes, this correlation is directly related to the spectral exponent α, allowing the dominant fluctuation type to be estimated from the time trace without explicitly calculating the power spectrum.

We calculate the spectral exponent *α* for each plateau and tethered bead and use the resulting *α* value to determine whether the uncertainty of the Allan variance or Hadamard variance can be calculated exactly using the analytical expressions derived by Greenhall and co-workers^58^. Specifically, exactu ncertainty estimates are only available within the ranges *α*_*AV*_ ∈[ −2,2]for the Allan variance and *α*_*HV*_ ∈[ −2,4] for the Hadamard variance. Traces that exhibit α values outside these ranges are typically dominated by strong low-frequency drift, high-frequency noise, or other nonstationary behavior, for which the magnetic trap stiffness estimation becomes unreliable.

Plateaus with α values within the corresponding validity ranges are classified as stable and kept for further analysis. In contrast, plateaus dominated by strong low-frequency drift or other nonstationary artifacts are excluded to avoid unreliable force and stiffness estimates.

### Determination of the Bead-Tether z-Offsets

The *z-offset* defines the origin of the z-axis and is, in turn, required for consistent force calibration. Although z-positions are measured on an absolute scale, the axis origin must still be defined. CALISTO follows the typical convention to set the bead’s z-position when it contacts the surface as *z*=0. Determining this contact position is essential for obtaining a meaningful force–extension relationship; skipping this step leads to unphysical extensions (including negative values) and incorrect force estimates, since the tether extension also enters Eq. (2).

Two commonly used approaches to determine the z-offset rely on defining conditions where the tether extension is effectively zero. One approach is to supercoil the DNA by applying rotations until the tether is fully coiled and the extension remains constant, which provides an accurate offset but requires torsionally constrained tethers and controlled magnet rotation. A more general and faster alternative, applicable also to non-coilable tethers, is to record bead motion at very low force (∼0 pN), where the bead fluctuates close to the surface and the extension is effectively zero. Measurements at high force (≥10 pN) define the (near) fully stretched state of the tether, while the low-force regime provides the surface-proximal reference point, enabling reliable determination of the bead–surface contact and thus the z-offset (Figure 3A). To determine the z-offset from a separate calibration dataset (time traces of all beads recorded at magnet positions corresponding to ≥10 pN, 0 pN, and again ≥10 pN), CALISTO provides three methods.

**Figure 3:**
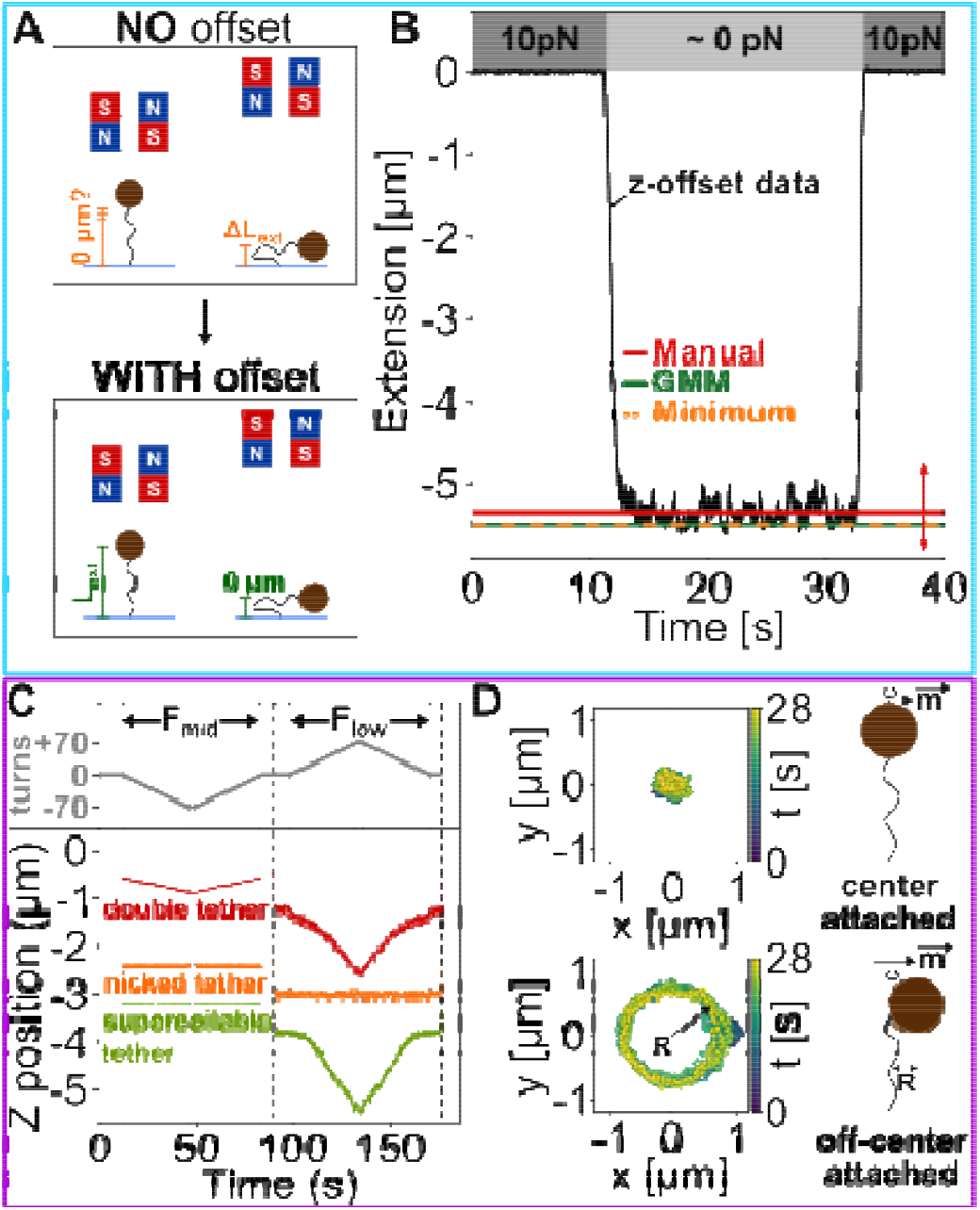
Z-offset determination and rotation-based tether classification. (A) Determination of the z-offset sets the zero point of the tether extension (L_ext_) axis to the extension where the bead makes contact with the surface, ensuring that the full DNA tether extension is represented as a positive value (lower panel). (B) Extension time trace of a DNA-tethered bead (M270, diameter: 2800 nm, black line) starting at a force of approximately 10 pN (Z_mag_ = 2.8 mm). The magnet is moved up to reduce the applied force to near 0 pN (Z_mag_ = 20 mm), such that the bead approaches the surface, before the magnet is lowered again and a high force is applied. Three methods are available for determining the offset from the test trace of the DNA-tethered bead: Gaussian mixture model (GMM, green), minimum value (orange) and a manual option (red). Red arrows illustrate the adjustability of the manual option using cursor input. (C) Rotation-based classification of DNA tethers into nicked DNA, supercoilable DNA, and multiple DNA tethers. Applying negative turns at high force (5 pN) decreases the extension in multiply tethered beads due to braid formation, while supercoilable and nicked tethers show no significant change. At low forces (0.5 pN), rotating the beads does not affect the extension of nicked tethers, as they are unable to form plectonemes unlike supercoilable tethers. (D) Tracked x,y-positions of a DNA tethered bead (M270, diameter: 2.8 m) while slowly applying -70 turns using the external magnets. x-y position traces reveal the attachment geometry. Beads tethered at the center exhibit a 2D Gaussian distribution, while off-center beads trace circular trajectories, with the radius reflecting the offset from the bead center.

A first simple option is the “Manual method”, where users simply inspect the trace and place a cursor at the desired position to set the z-offset. Second, a straight-forward automatic approach selects the “minimum z-position” in the dataset after smoothing the data using a third-order Savitzky–Golay filter. Since the filter window length is a free parameter, we determine it from the Allan variance of the trace. The minimum of the AV corresponds to the averaging window that minimizes fluctuations between consecutive averaged data points. We then use half of this window length for the Savitzky-Golay filter, ensuring, according to the Nyquist criterion, that features occurring on the timescale of the AV minimum remain resolved while noise is reduced.

A third alternative approach for the z-offset determination is to apply a Gaussian mixture model (GMM). This method estimates the minimum value of a given trace using GMMs with varying configurations. A GMM with *K* components is defined by the probability density function

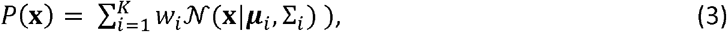

where *w*_*i*_ represents the weight of the *i*-th Gaussian component with Σ*w*_*i*_=1, and *N* (**x**| *μ*_*i*_, Σ_*i*_) is the normal distribution with mean vector μ_*i*_ and covariance matrix *Σ*_*i*_. We fit two GMM with 4 and 6 components to the extension values of the given trace to separate the stationary and transient regions where the GMM with 6 components represents a finer separation. The z-offset trace shows a trapezoidal shape (Figure 3B) with three plateaus and two edges, resulting in five regions in total. For fast edges, their contribution can be neglected, leaving three distinct regions. We therefore use 4 or 6 GMM components, including an extra component to flexibly capture the trapezoidal shape while remaining robust to noise. Next, we identify the component with the lowest z component of the mean across both GMMs and estimate the z-offset by subtracting one standard deviation from the mean.

We compare and validate the different methods for z-offset determination by comparing it with the results from fully coiling the DNA tethers via magnet rotation (Supporting Information Section S1). In the rotation assay, increasing the number of magnet turns introduces supercoils that progressively shorten the tether. When the bead contacts the surface, the extension reaches a constant value, which is taken as the offset. The offsets, which were obtained from traces at very low forces using the minimum and GMM approaches, closely match the surface-contact extension determined by supercoiling the DNA tethers. This finding demonstrates that the methods reliably estimate the zero-offset without active rotation. Importantly, unlike rotation-based approaches, the minimum and GMM methods can also be applied to non-supercoilable tethers, making it suitable for assays where torsional manipulation is not possible.

### Closed-Form Expressions for the Power Spectral Density, Allan Variance, and Hadamard Variance

Determining the trap stiffness (*κ*) is an integral part of the force calibration since it, together with the average extension, relates the transverse fluctuations of the bead to the applied force. Note that, we use *κ* instead of the previously used *κ*|| for brevity. In the experiment, the bead continuously fluctuates because of thermal collisions with molecules in the surrounding buffer. The magnitude and timescale of these fluctuations depend on both the trap stiffness and the viscous drag acting on the bead. We therefore analyze the measured bead-position time traces to determine (*κ*) and the drag coefficient *γ*). We model the motion of the bead-tether system in the axis parallel to the magnetic field as a harmonic trap of stiffness *κ* centered at force-balance position. Random collisions between the bead and the surrounding buffer are represented by a stochastic Langevin force *FL*, which obeys the fluctuation-dissipation relation, ⟨*FL*(*t*)*FL*(*t*′)⟩ = 2*γkBTδ*(*t* − *t*′), where *γ* is the viscous drag coefficient, and *δ*(*t* − *t*′) is the Dirac delta distribution. Since inertial effects are negligible on the experimentally relevant timescales, the bead motion is well-described by the overdamped Langevin equation

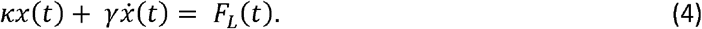

In order to extract the trap stiffness *κ* and drag coefficient *γ* from the experimental bead traces, we use the closed-form expressions of the PSD, AV, and HV for the overdamped Langevin equation (Eq. (4)) described previously^48, 51, 52^. The PSD and AV methods are commonly used to estimate forces from tweezer time traces but can be affected by drift and other measurement imperfections. In contrast, the Hadamard variance is robust against drift, especially linear drift.

#### Power Spectral Density (PSD)

The downsampling-corrected analytical closed-form expression of the PSD is given as^48, 52^

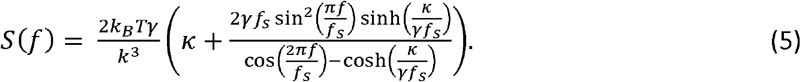

With *f*_*s*_ as the sampling frequency. Eq. (5) specifically applies when the camera exposure time *τ*_0_ equals the sampling time *τ*_*s*_ = 1/*f*_*s*_,i.e. *τ*_*0*_ = *τ*_*s*_. This assumption holds for most modern CMOS cameras since their dead time is negligible. Eq. (5) is fit to the PSD that is computed from the position time traces to determine *k* and *γ*.

#### Allan Variance (AV)

The closed-form expression of the AV for Eq. (5) is then given by^48^

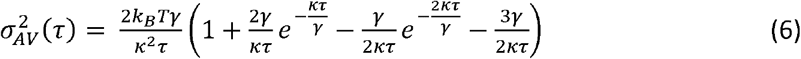

with *τ* being the observation time intervals. Hence, in order to use the AV to obtain *K*, we calculate the AV of the measured trajectory of the bead. We then fit the calculated AV to Eq. (6) with *K* and *γ* being fitting parameters using maximum likelihood estimation (MLE).

#### Hadamard Variance (HV)

The closed-form expression of the HV of the bead motion then follows as^51^

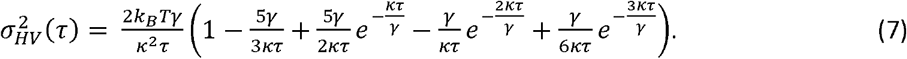

To determine the trap stiffness and drag coefficient from the HV curves, we again employ an MLE fit of the HV as a function of observation times.

In principle, fluctuations perpendicular to both the axial force (*F*_*z*_) and the magnetic field could be used for force calibration. However, this approach requires correcting the measured molecular extension for bead rotation in the magnetic field^50^ (i.e. replacing <*L*_ext_ > in Eq. (2) by (<*L*_ext_ > + *R*_bead_ )), a correction not implemented in CALISTO. Therefore, calibration should be performed along the transverse direction that is parallel to the magnetic field. Furthermore, CALISTO employs a skewed Gaussian distribution to extract the z position and tether extension. The skewed distribution is a generalized Gaussian form that accounts for surface damping effects that arise due to the interactions between surface and magnetic bead.^59^

### Worm-Like Chain Model

One way to assess the integrity of a DNA-tethered bead is to determine whether its force-extension behavior is consistent with that expected for DNA. At forces up to approximately 10 pN, double-stranded DNA is well described by the inextensible worm-like chain (WLC) model.

The WLC model characterizes the DNA using two parameters: the contour length *L*_*C*_, which is the total length of the DNA molecule, and the persistence length *L*_*p*_, which describes the distance over which the DNA orientation remains correlated. The expected contour length depends primarily on the length of the DNA construct and can be estimated from the number of base pairs using approximately 0.34 nm per base pair.^39^ The persistence length is independent of the total DNA length but depends on experimental conditions, particularly the ionic strength of the buffer, and is typically approximately 40-50 nm under commonly used aqueous buffer conditions.^59-61^

To determine*L*_*p*_ and *L*_*C*_ from the force-extension data measured in the magnetic-tweezers experiment, we use a high-accuracy interpolation of the inextensible WLC model described in Ref. ^40^:

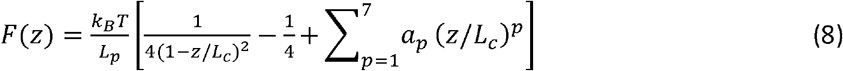

where *F* is the applied force, *z* is the measured DNA extension, *k*_*B*_ is the Boltzmann constant, and *T* is the absolute temperature. The coefficients are{a1‖,}={1,−0.5164228, −2.737418,16.07497, −38.87607,39.49944, −14.17718}.

These coefficients are fixed values taken from Ref. and are not determined separately for each experimental dataset. They define a polynomial correction to the commonly used Marko-Siggia approximation. Thus, only *L*_*p*_ and *L*_*c*_ are varied when fitting the experimental force-extension data.

Since the inextensible WLC model provides a reliable description of double-stranded DNA only at relatively low forces, the fit is restricted to forces below approximately 10 pN. The resulting values of *L*_*p*_ and *L*_*c*_ can then be compared with the values expected for the DNA construct and buffer conditions by the user. Substantial deviations may indicate, for example, an incorrect tether length, multiple DNA tethers, nonspecific interactions, or another compromised tether configuration. Such beads can therefore be identified as unsuitable for inclusion in the force-calibration dataset.

For visualization in the graphical user interface, the fitted curve is extrapolated beyond the fitting range to the maximum experimentally measured force. This extrapolated part of the curve is displayed only as a visual guide and is not used to determine *L*_*p*_ or *L*_*c*_.

### Exponential Fit Functions for Force vs Magnet Position Curves

To provide a convenient empirical description of the force–magnet position relationship, we fit the data with a two-component exponential model. Compared with a single exponential, the additional exponential term substantially improved the description of the measured force curves while retaining a simple analytical form. The model can also naturally reduce to a single exponential when one component dominates (i.e., *c* → 0 or 1, or *λ*_1_ ≈ *λ*_*s*_), avoiding unnecessary model complexity when the data do not support it. The force–magnet position curves were fitted using

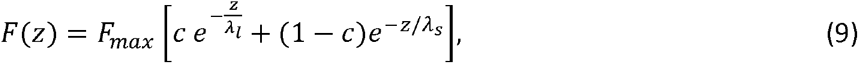

with the constraints *F*_max_ ≥ 0, *λ*_1_> *λ*_*s*_ and *c* ∈ [0, 1]. Because the measurement uncertainty increases approximately in proportion to the force, resulting in heteroscedastic (multiplicative) noise, parameter estimation was performed using an iterative reweighted least-squares (IRLS) approach^51^(Supporting Information Section S2). For fixed length scales (*λ*_1_, *λ*_*s*_), the amplitude and mixing term (*F*_max_, *c*) were determined using non-negative least squares. Optimization of the length scales was carried out using the L-BFGS-B algorithm^62^−^66^. The model was fit using multiple random initializations to avoid local minima.

### Statistical Tests

To quantitatively evaluate the different stages of the calibration pipeline, including reference-bead selection, optimization of magnet-position parsing parameters, and filtering of stable force plateaus, we compared the measured forces with the fitted double-exponential calibration model using complementary statistical metrics. RMSE was used to quantify the absolute deviation between measured and predicted forces,

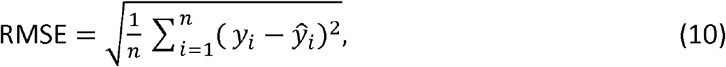

whereas the reduced chi-squared statistic,

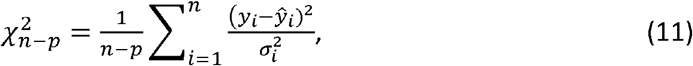

additionally accounts for the estimated measurement uncertainties. Here, is the number of observations, *p* the number of fitted parameters, and 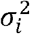 the estimated uncertainty of each force measurement. Cross-validated RMSE was additionally used to assess predictive performance on data not included in the respective model fit, thereby distinguishing improvements in generalizability from improvements in the fit to the calibration data alone.

For plateau filtering (see *Noise Classification above*), we additionally tested whether the observed changes in calibration performance could be attributed solely to the reduced number of data points or whether filtering systematically altered the underlying force–magnet position relationship. First, a size-matched subsampling test was performed by repeatedly drawing random subsets from the unfiltered dataset with the same number of observations as the filtered dataset and comparing their calibration metrics with those obtained after filtering. This provides the expected variation resulting from dataset-size reduction alone. Second, a permutation test was used to test whether the filtered and unfiltered datasets were statistically distinguishable beyond random variation by repeatedly permuting their group assignments and comparing the resulting differences in calibration metrics with the observed differences. Empirical p-values were calculated as the fraction of randomized datasets yielding differences equal to or larger than those observed experimentally.

In the following three sections, we describe the magnetic-tweezers setup and force-calibration measurements, as well as the preparation of DNA and flow-cell samples used in this study to test the CALISTO pipeline.

### Magnetic Tweezer Setup

We utilize a custom-built MT setup equipped with two linear stepper motors (M-126.PD2 and C-150.PD, Physik Instrumente, Germany) to control the height and rotation of two external NdFeP permanent magnets (size: 5×5×5 mm^3^, N50 grade, 1 mm gap, Supermagnete, Germany) above the flow cell.^17, 67-69^ Additionally, a Pifoc piezo stage (PifocP726.1CD; Physik Instrumente, Germany) adjusts the focus of the objective (40x oil immersion, Olympus, Japan). Imaging of the diffraction rings of the magnetic beads is performed using a tube lens (*f*= 150 mm), a red LED (69647; LUMITRONIX LED Technik, Germany) for illumination, and a CMOS camera (12M Falcon2; Teledyne DALSA, Waterloo, Canada) recording at 58 Hz for the full field of view. For some measurement we reduce the field of view and record at 160 Hz. Bead tracking is conducted using an open-source LabVIEW code, which employs a CUDA-based parallel computing algorithm and quadrant interpolation.^24, 45^

### DNA and Flow Cell Preparation

We use a 21 kbp DNA (dsDNA) construct derived from lambda phage DNA, prepared as previously described.^70^ The terminal regions (∼600 bp) are functionalized with multiple digoxigenin (Roche, The Netherlands) moieties at one end and biotin (Sigma Aldrich, USA) moieties at the other end, respectively, for attachment. Flow cells are constructed using two glass coverslips. The bottom slide (24 × 60 mm^2^, #1, VWR, Germany) is first functionalized with (3-Glycidoxypropyl)-trimethoxysilane (Thermo Fisher Scientific, USA) and then incubated with 1 or 3 µm polystyrene beads (Polysciences, USA) in ethanol (VWR, USA) to serve as reference beads for drift correction. The top slide (24 × 60 mm^2^, #1.5, Carl Roth, Germany) has two ∼1 mm diameter holes to connect to the pump system (Ismatec, Germany) for fluid exchange in the flow chamber. Both slides sandwich a single layer of melted parafilm (Carl Roth, Germany) with a cutout, forming a ∼25 µl channel that connects the inlet and outlet of the flow chamber. After assembly into a custom-built flow cell holder, the flow cell is incubated with 100 µg/ml anti-digoxigenin (Abcam, UK) in 1x PBS buffer (Sigma Aldrich, USA) for at least 2 hours to facilitate DNA attachment via digoxigenin–anti-digoxigenin binding. The channel is then rinsed with 1x PBS and passivated with 100 mg/ml BSA (Carl Roth, Germany) for 1 hour to minimize non-specific interactions, followed by another PBS rinse. We use commercially available streptavidin-coated magnetic beads, specifically 1 µm DynaBeads MyOne Streptavidin C1 and 2.8 µm DynaBeads M-270 Streptavidin (Thermo Fisher Scientific, USA), to attach to the DNA via streptavidin-biotin interaction. To couple the beads, 2 µl of picomolar DNA stock is mixed with either 2 µl of MyOne beads or 13 µl of M-270 beads in 100 µl of 1x PBS for approximately 30 s. The bead mixture is then added to the flow cell and incubated for several minutes before unbound material is flushed out with 1x PBS.

### Force Calibration Measurement

For force calibration measurements, we first determine the distance between the magnet and the top coverslip of the flow cell by gradually lowering the magnet until it gently touches the top surface. The magnet position origin is defined as the point where the magnets contact the top coverslip. To establish the bead–surface coordinate system, a separate z-offset calibration dataset is recorded by acquiring time traces of all beads at three magnet positions corresponding to high force (≥10 pN), very low force (∼0 pN), and again high force (≥10 pN). Next, tethered magnetic beads are screened to identify those attached via multiple DNA molecules by analyzing their response to applied force and torque. We first apply negative rotations (−70 turns) at intermediate forces, i.e. ∼ 5 pN, where no change in extension is expected for beads tethered by a single double-stranded DNA molecule. Beads attached via multiple tethers form braids under negative rotation, leading to a reduction in extension; these beads are excluded from measurements. Further screening is conducted to identify nicked DNA tethers by overwinding the DNA at low force. We apply positive rotations (70 turns) at low forces, i.e. ∼0.4 pN, and record the tether extension. Nicked tethers, unable to form plectonemes under low force, exhibit no change in extension. For the force calibration measurement, we record time traces of the bead-tether complex at 49 different magnet positions at a temperature of 293 K.

## USER MANUAL AND MAIN FINDINGS

CALISTO is an open-source software for analyzing magnetic tweezer (MT) force calibration experiments. It was developed for thermal-motion-based force calibration from bead position time traces by combining automated analysis routines with an interactive graphical user interface (GUI). The software provides a structured and intuitive workflow for processing, inspecting, and refining experimental data. CALISTO is designed for single-molecule force spectroscopy (SMFS) experiments, including high-throughput measurements in which many DNA-tethered beads are tracked simultaneously. It provides a standardized yet flexible framework for force calibration, improving the consistency and reproducibility of data analysis across different experimental setups.

In the following sections, we provide an overview of the main functionalities of CALISTO and describe the individual analysis modules through a detailed user guide. We also introduce several new analysis features that are not part of standard magnetic tweezer calibration workflows. These additions improve the accuracy, robustness, and reliability of force calibration, particularly for large datasets and long measurements. The graphical user interface includes tools for convenient data visualization and exploration. Users can display traces on linear or logarithmic axes, apply adjustable downsampling for faster rendering of large datasets, and focus on selected time intervals or traces of interest. All plots can be exported either as high-quality image files (PNG, JPEG, or SVG) or as raw numerical data in CSV or HDF5 format for further analysis in external software.

### Main Functionalities of CALISTO

The first step in force calibration is loading the experimental data and defining the basic measurement settings. CALISTO simplifies this process by storing setup-specific parameters internally in a YAML file with custom setup-name, which can be edited by the user. Users then select their magnetic-tweezers setup from a dropdown menu, after which the corresponding parameters, such as unit conversions, scaling factors, correction values, and the variance axis used for fluctuation analysis, are applied automatically.

In magnetic tweezers, the bead undergoes thermal fluctuations in all spatial directions. CALISTO analyzes the fluctuations along the magnetic field direction, typically chosen based on the magnet geometry. Although fluctuations along other directions can be used, doing so would require additional assumptions about bead rotation and the full bead-motion model in the magnetic field.^50^ To avoid introducing these extra model assumptions, CALISTO currently restricts the fluctuation analysis to the selected variance axis.

CALISTO supports both plain text and HDF5 input formats, which can be selected through a drop-down menu. For plain text input, two separate files must be loaded: one containing the bead position traces (of reference and DNA-tethered beads) and another containing the motor movement data. For HDF5 input, bead traces and magnet movement must be stored in a single file structure. Additional information about supported formats and file organization, and a detailed user manual and link to sample data are available in the GitHub repository (github.com/lipfertlab/calisto).

Before starting the analysis, several global experimental parameters must be defined. For example, the sampling frequency, i.e. how many images are recorded per second, is automatically extracted from the raw data and displayed in the interface, but it can also be adjusted manually if needed.

Accurate force calibration requires that the camera records images fast enough to resolve the thermal motion of the bead–tether system.^47, 50^ In a magnetic tweezer experiment, the bead constantly undergoes thermal fluctuations. At the same time, the DNA tether and the applied magnetic force confine the bead around an average position, modelled as a harmonically trapped particle undergoing Brownian dynamics.

The properties of these fluctuations depend on several experimental parameters, including the bead size, the DNA length, and the applied force. Larger beads, shorter DNA tethers, and higher forces generally lead to more confined fluctuations (Supporting Information Section S3). These fluctuations are characterized by a characteristic time *τ*_*c*_, which describes how quickly the bead position loses memory of its previous state. To accurately capture the thermal fluctuation, the camera frame rate must be sufficiently high. If the frame rate is too low, fast fluctuations are missed and the camera acts as a low-pass filter, which reduces the measured variance of the fluctuations and leads to a systematic overestimation of the applied force. To help users choose suitable acquisition settings, we provide a Python script that estimates the characteristic fluctuation frequency and the corresponding minimum frame rate based on the experimental parameters and force range (Supporting Information Section S3).

In addition to the frame rate, the recording time at each magnet position is important. Reliable force calibration requires sampling of the fluctuations sufficiently long. The relative statistical uncertainty approximately scales as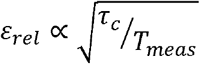, where *τ*_*c*_ is the characteristic fluctuation time and ***T***_*meas*_ is the total measurement time.^71^ This relation shows that reducing the uncertainty requires recording substantially longer than the intrinsic fluctuation time of the system. For example, achieving a relative uncertainty of approximately 10% requires a measurement time roughly 100 times longer than *τ*_*c*_. Consequently, low-force measurements, which have longer characteristic times, require longer measurement times than high-force measurements. We therefore also provide a script that estimates the minimum recommended recording time to achieve a set number of independent datapoints (Supporting Information Section S3). For example, using M270 beads at a force of 10 pN, the recommended measurement time is > 4 s for a 5 kbp DNA tether, whereas it increases to > 10 s for a 21 kbp tether.

Several additional experimental parameters are required for force calibration. The system temperature is needed for calculations involving the power spectral density (PSD), Allan variance (AV), Hadamard variance (HV), and worm-like chain (WLC) models.

The bead radius is used only as an initial estimate for the drag coefficient during fitting. The final fitted drag coefficient is determined directly from the experimental data, making the calibration relatively insensitive to the user-provided bead size. As a result, approximate manufacturer specifications are generally sufficient, even when the precise bead diameter is not known accurately.

If specified in the configuration file, the variance axis is loaded automatically but can still be adjusted manually.

Once all required inputs have been provided, the “Start Force Calibration Session” button becomes active. If any required information is missing, the button remains inactive to prevent incomplete analysis. Starting a calibration session opens CALISTO’s main analysis window, which contains five panels: “Check Data,” “Offsets,” “Rotation Test,” “Force Calibration,” and a bead property table.

The bead property table provides a compact overview of the analysis status for each bead and helps users identify suitable calibration traces. It summarizes the bead type (reference bead or DNA-tethered magnetic bead), tether type (supercoilable, nicked, or multiple-attached), offset values, quality control results, and inclusion status for the final calibration curve. Beads excluded during the workflow are labeled together with the corresponding exclusion reason. The individual panels and analysis steps are described in detail in the following sections.

### Check Data Panel

The “Check Data” panel is the starting point for data quality control. It allows users to inspect, classify, and organize the position traces of both reference beads and DNA-tethered beads. After opening the “Choose and Plot Traces” window, users can display one or more traces simultaneously and assess their quality before proceeding with force calibration.

Reference beads are evaluated first because they are required for mechanical drift correction. Without reliable reference beads, slow movements of the microscope, sample stage, or fluid chamber cannot be removed from the data. Since drift correction affects all subsequent analysis steps, identifying suitable reference beads is a prerequisite for accurate force calibration.

A high-quality reference bead exhibits stable fluctuations around a constant mean position and shows no jumps, or tracking artifacts. In contrast, low-quality traces often contain sudden position shifts, intervals of drift, unusually large fluctuations, or inconsistent behavior across the x-, y-, and z-axes. Such traces should be excluded because they can distort the drift correction and introduce systematic errors into the force calibration.

Users can manually classify traces as “high-quality” or “low-quality” based on visual inspection. To make this process more objective and reproducible, CALISTO additionally provides a semi-automated reference bead selection algorithm that can be activated via the “Identify Suitable Reference Beads” button. Rather than relying on a single threshold, the algorithm identifies traces whose statistical properties are consistent with a common trend. In particular, traces are smoothed, compared against the ensemble of reference traces, and screened for outliers based on their noise level, drift behavior, and deviation from the average trend (see Materials and Methods for details). Beads exhibiting unusually large fluctuations or systematic deviations are flagged for exclusion. In addition, users can inspect the resulting selections for each spatial axis individually and manually modify them if necessary. This combination of automated screening and user supervision helps ensure that the final calibration is based on a consistent set of high-quality reference traces.

Figure 2A shows representative examples of high- and low-quality reference bead traces. To provide a practical benchmark for expected noise levels, we characterized the tracking precision using surface-immobilized beads under stable experimental conditions (Supporting Information Section S4). For 1 µm beads, the standard deviation ranges from approximately 0.5 to 4 nm depending on the measurement axis and acquisition frequency. These values provide a useful rule of thumb when assessing data quality. Significantly larger fluctuations may indicate tracking problems, poor illumination, or mechanical instability.^46^ As a rough estimate, the scattered light intensity goes as ∼ R_bead_^2^ (since the ∼µm-sized beads predominantly scatter in the Mie regime), i.e. imaging a 3 µm bead yields 9 times as many photons as imaging a 1 µm bead. Since the standard deviation *σ* scales inversely with the number of photons, i.e. 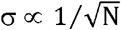, increasing the bead diameter from 1 µm to 3 µm would therefore reduce the standard deviation by approximately a factor of 3.^46^

After suitable reference beads have been identified, users can evaluate the DNA-tethered bead traces. High-quality tether traces exhibit stable fluctuations around a well-defined mean position and remain free of obvious tracking artifacts. In contrast, low-quality traces frequently contain sharp spikes in the z-position signal, sudden jumps, intermittent loss of tracking, or other nonphysical features.

Although isolated spikes do not necessarily change the mean position if they occur symmetrically around the average, they are generally indicative of tracking errors. For this reason, traces containing frequent or pronounced spikes should typically be excluded from force calibration. Such artifacts reduce confidence in the measured bead position and may bias both the variance of the tether extension and, therefore, the calculated force. Figure 2A shows representative examples of acceptable and problematic tether traces.

To identify regions corresponding to constant force, the “Plot and Partition Magnet Positions” module analyzes the recorded magnet position data. This procedure consists of two steps. First, the software distinguishes between periods in which the magnets are moving and periods in which they remain stationary. Consecutive motor positions differing by less than a user-defined threshold are classified as stationary. Second, users can define a minimum plateau duration to exclude short segments that are unlikely to represent true equilibrium measurements. Since the partitioning is performed automatically by the analysis script, users should verify that the detected plateaus correspond to the expected stationary magnet positions before proceeding with further analysis.

Careful selection of these parameters is important. If the threshold is too permissive or the minimum plateau duration too short, transient motor movements may be misclassified as stable force plateaus. Such short regions can arise from motor hysteresis, small positioning steps, or delayed motor responses and should generally be excluded from calibration (Figure 2B).^72, 73^

The “Identify Suitable Reference Beads” module further assists drift correction through a semi-automated workflow consisting of four steps: trace smoothing, outlier detection, extraction of the mean drift trend, and final refinement of the reference bead set.

### Offsets Panel

The “Offsets” panel is used to determine the zero position of the z-axis for each bead, which is essential for obtaining meaningful force and extension measurements. In magnetic tweezers, the z-position is measured on an absolute scale because vertical movement of the microscope objective is calibrated and corrections for the oil–water interface are known. However, the choice of where z = 0 is located remains arbitrary. CALISTO adopts the common convention that z=0 corresponds to the bead’s z position when it contacts the surface (Figure 3A).

Determining this contact position, referred to here as the “offset,” is critical for calculating the correct tether extension. If the offset is not determined properly, the calculated tether length can become unphysical, including negative extensions, which directly leads to incorrect force estimates through the extension term in Eq. (2).

The software provides three approaches for determining offsets. First, users can extract offsets directly from dedicated offset traces recorded at both very low and high forces. These traces are analyzed bead by bead using one of three methods: a Gaussian Mixture Model (GMM), a minimum-value approach, or a fully manual mode in which users select the offset position interactively using the cursor. Details about the minimum and GMM methods can be found in the Materials and Methods section.

The manual option is particularly important because experimental traces can vary substantially between setups and measurement protocols. For example, some experiments contain only two force levels, whereas others may contain several intermediate force steps or more complex bead behavior. Manual selection therefore provides a fallback option whenever automated methods fail or produce ambiguous results.

Among the automated approaches, the minimum-value method is conceptually the simplest and likely the most broadly applicable. After smoothing the trace to suppress short-time fluctuations, the minimum of the smoothed trace is interpreted as the bead position at surface contact. This approach works well for clean datasets and does not rely on assumptions about the detailed shape of the position distribution.

However, the minimum-value method can be sensitive to tracking artifacts or isolated outliers. Erroneous tracking points can artificially shift the detected minimum and therefore bias the offset estimate. The GMM-based approach is generally more robust against such artifacts because it analyzes the full distribution of bead positions rather than relying on a single extreme value. In practice, the GMM separates the trace into different statistical clusters, allowing the contact position to be identified more reliably even when occasional tracking errors or spikes are present. For this reason, CALISTO uses the GMM approach as the default automated method, while still providing the minimum-value method as a simpler and more universal alternative.

If an offset analysis has already been performed previously, users can directly load an existing table of offset values. Alternatively, if no offset traces are available, a constant offset value can be entered manually.

To ensure that only physically meaningful tethers are included in the analysis, users can define an acceptable offset range. Beads with offsets outside this interval are automatically excluded. Extremely small offsets may indicate beads stuck close to the surface or incorrectly tracked particles, whereas unusually large offsets can indicate multiple tethers, tracking errors, or incorrect bead identification.

### Rotation Test Panel

The “Rotation Test” panel helps users determine the tether type and evaluate how the bead is attached by analyzing rotation-test measurements. All panels are synchronized, allowing users to inspect the same time intervals across different data representations simultaneously.

After loading the relevant files, the software displays the z-position trace of the selected bead during controlled bead rotation. The response of the tether to rotation reveals important information about the tether geometry and topology.

As shown in Figure 3C, applying negative rotations to underwind DNA at intermediate force shortens multiple-tethered beads because the two DNA molecules wrap around each other and form a braid. In contrast, supercoilable and nicked single tethers show little or no shortening under the same conditions. At low forces, positive rotations to overwind DNA further distinguish supercoilable from nicked tethers: supercoilable DNA forms plectonemes and shortens, whereas nicked DNA cannot store torsional stress and the extension therefore remains unchanged.

In addition to the z-position traces, the software displays the x-y motion of the bead during rotation (Figure 3D). A bead attached close to its center fluctuates symmetrically and produces an approximately 2D Gaussian distribution. In contrast, an off-center tether causes the bead to rotate along a circular trajectory. The radius of this circle directly reflects the distance between the tether attachment point and the bead center.^49, 74^

Users can inspect individual beads manually to classify tether types and determine whether the attachment geometry is suitable for force calibration. Multiple tethers should generally be excluded because they distort the force–extension relation.

Off-center tether attachment can also introduce systematic force errors, particularly when the tether length becomes comparable to the bead radius. In this regime, geometric effects modify the effective extension of the tether and therefore bias the force estimate. For long DNA tethers and small beads, however, this effect becomes relatively small.

For larger beads such as 2.8 µm M270 beads, we recommend excluding beads with off-center attachment distances larger than approximately 500 nm. This threshold was estimated geometrically for 21 kbp DNA tethers and corresponds to a force error exceeding roughly 2% (Supporting Information Section S5). The exact value depends on bead size and tether length, but 500 nm provides a practical guideline that balances calibration accuracy against retaining sufficient bead statistics.

### Force Calibration Panel

The “Force Calibration” panel allows users to inspect force calibration results for individual beads and generate the final calibration curve using only traces that pass all quality-control steps (Figure 4). The “get forces per bead” panel displays both the force–magnet position relation and the force–extension behavior per tether, enabling users to identify problematic measurements before constructing the final master calibration curve.

**Figure 4:**
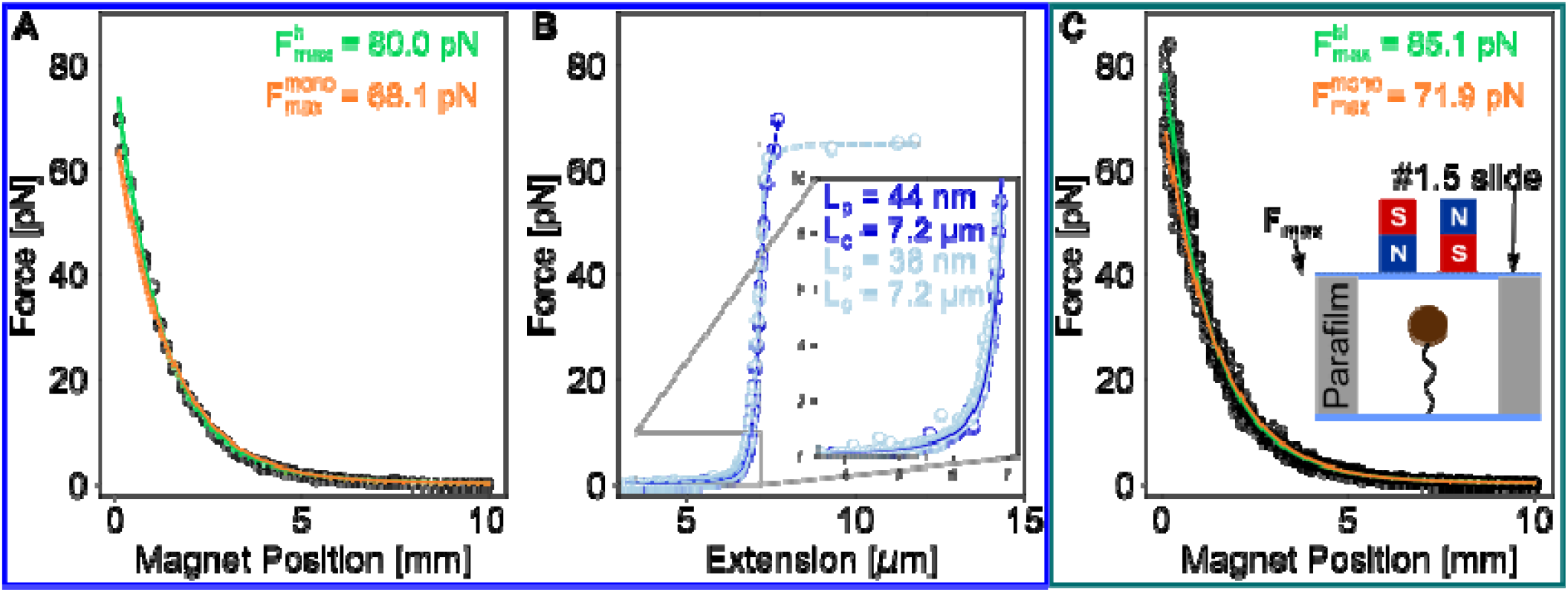
Force vs. magnet height and DNA force-extension relations. (A) Force versus magnet position plot (based on HV analysis) for a single M270 (diameter: 2.8 m) bead tethered to a 21 kbp dsDNA. A single-exponential fit provides an approximate fit (orange line, ), but the data are more accurately described by a bi-exponential fit (green line; ), which yields a maximum force of 80.0 pN. (B) HV-based force–extension plot for a supercoilable dsDNA tether (dark blue) and a nicked dsDNA tether that shows the characteristic overstretching transition at ∼65 pN (light blue). Inset: Worm-like chain (WLC) fit to both data curves gives a contour length of 7.2 µm and a persistence length of 45 nm (dark blue) and 38 nm (light blue) for the supercoilable (dark blue) and overstretching (light blue) tether, respectively. (C) The final master force calibration curve is derived by fitting all data points from 30 different beads. The bi-exponential fit (green line) indicates a maximum force of 85.1 pN. The fit parameters are summarized in Table 1.

CALISTO implements both mono-exponential and bi-exponential fitting models for the force– magnet position calibration, as both approaches have been widely used in the literature and provide a good description of the force dependence on magnet position for typical magnet geometries employed in MT. ^1, 75, 76^ To help users evaluate which model better describes the data, the software additionally reports the Akaike Information Criterion (AIC)^77^. In general, a lower AIC value indicates a better compromise between fitting accuracy and model complexity, and therefore suggests a more appropriate model for the experimental data.

**Table 1:** Comparison of force–magnet distance calibration curves. obtained in this work with previously published magnetic tweezer calibrations from Lipfert et al. (PNAS, 2014) and Yu et al. (RSI, 2014) for M270 and MyOne magnetic beads. The table summarizes the maximum force F_max_, the fitting model used for calibration, and the corresponding analytical force–distance relation. The maximum forces are interpolated to the contact point between upper cover glass and external magnet accounting for the different flowcell geometries (cover glass thickness - #1: D_#1_=0.15mm, #1.5: D_#1.5_=0.18 mm; and number of parafilm layers -D_OneParafilm_=0.13 mm)).

| Bead Type | Study | $F_{\max}$ (pN) | Calibration model | Calibration curve |
| --- | --- | --- | --- | --- |
| M270 | This work | 85.1 | Weighted bi-exponential | $F(z) = 85.13\text{ pN} \cdot (0.50 e^{-z/0.92\text{ mm}} + (1-0.50) e^{-z/1.63\text{ mm}})$ |
| M270 | Lipfert et al. PNAS (2014) | 83.3 | Mono-exponential | $F(z) = 76.83\text{ pN} e^{-z/1.36\text{ mm}} + 0.64\text{ pN}$ |
| M270 | Yu et al, RSI (2014) | 78.7 | Bi-exponential with offset | $F(z) = -2.74\text{ pN} + 71.8\text{ pN} \cdot e^{(-z/1.33\text{ mm})} + 4.0\text{ pN} e^{(-z/36.9\text{ mm})}$ |
| MyOne | This work | 10.4<br>12.6 | Weighted bi-exponential | $F(z) = 10.41\text{ pN} \cdot (0.58 e^{-z/0.89\text{ mm}} + (1-0.58) e^{-z/1.63\text{ mm}})$<br>$F(z) = 12.55\text{ pN} \cdot (0.82 e^{-z/1.06\text{ mm}} + (1-0.82) e^{-z/2.39\text{ mm}})$ |
| MyOne | Lipfert et al. PNAS (2014) | 9.47 | Bi-exponential | $F(z) = 8.0\text{ pN} e^{-z/1.23\text{ mm}} + 0.76\text{ pN} e^{-z/2.48\text{ mm}}$ |
| MyOne | Yu et al, RSI (2014) | 9.49 | Bi-exponential with offset | $F(z) = -0.117\text{ pN} + 8.68\text{ pN} \cdot e^{(-z/1.25\text{ mm})} + 0.2\text{ pN} e^{(-z/14.5\text{ mm})}$ |

Careful inspection of individual tethered beads is important because not all traces behave physically as expected. For example, some beads show force–magnet position curves that deviate strongly from the expected exponential behavior. Similarly, force–extension curves can yield unrealistic contour lengths or persistence lengths when fitted with the worm-like chain (WLC) model.^40^ Such deviations are often caused by tracking artifacts, multiple DNA attachments, damaged tethers, or incorrect bead attachment geometries.

Figure 4A shows a representative force–magnet position calibration obtained from a single M270 bead tethered to 21 kbp DNA over a force range extending to 80 pN. The data are fitted using a weighted bi-exponential model described in the Materials and Methods section.

Importantly, the variability in force calibration measurements is not constant across the full force range. At higher forces, the fluctuations and fitting residuals become larger in absolute magnitude since bead-to-bead variations cause the variance of the force to scale as F^2^ (as, in particular, bead-to-bead variation in size and magnetic content lead to an approximately constant relative force error), which leads to so-called heteroscedastic noise. To account for this behavior, CALISTO implements weighted fitting using iterative reweighted least squares (IRLS) with weights proportional to 1/F^2^. This weighting prevents the high-force data points from artificially dominating the fit and improves the overall calibration accuracy across the entire magnet range.

We evaluated the different fitting approaches using calibration data from 26 independent M270 beads (Supporting Information Section S6). While unweighted mono-and bi-exponential fits already perform reasonably well, the weighted bi-exponential approach consistently reduced the residual spread and improved the information criteria. These results show that proper weighting becomes particularly important when fitting over a broad force range that includes high forces approaching or even exceeding 80 pN.

Figure 4B shows representative force–extension curves for two different tether types: a supercoilable DNA tether and a nicked tether. The nicked tether exhibits the characteristic overstretching transition near 65 pN, in agreement with established literature values.^42^ In both cases, the fitted contour length corresponds closely to the expected crystallographic DNA length for 21 kbp (7.14 µm). For the supercoilable tether, the fitted persistence length falls within the expected range of approximately 40–50 nm.^59^ Beads that produce unrealistic contour lengths, implausible persistence lengths, or abnormal overstretching behavior should be excluded from the final calibration dataset by the user.

Figure 4C shows the resulting master calibration curve generated from 30 independent DNA tethers that passed all quality criteria, including the absence of overstretching artifacts and off-center attachment distances larger than 500 nm.

The bi-exponential model again provided a substantially better description of the final calibration curve than the mono-exponential model, as reflected by the lower AIC value (AIC_bi = −1158.10 versus AIC__mono = −862.06).

For the WLC analysis, we acquired force–extension data at approximately 30–40 discrete magnet positions spanning forces between 0 and 10 pN. This dense sampling improves the robustness of the contour-length and persistence-length fits. In practice, fewer magnet positions can already be sufficient, provided that the full force range is sampled adequately and the force–extension curve remains well constrained.

To facilitate comparison between different magnetic tweezer setups, we summarize the fitted calibration parameters in Table 1. The resulting force–distance relations agree well with previously published calibrations. For example, after accounting for differences in flow-cell geometry, cover-glass thickness, and parafilm spacing, the extrapolated forces at zero magnet distance are fully consistent with earlier reports from Lipfert et al.^34^ and Yu et al^75^. In particular, both previous studies reported forces near 10 pN for MyOne beads and approximately 80–90 pN for M270 beads at minimal magnet distances, which agrees closely with the values obtained here.

To further validate CALISTO, we additionally performed calibrations using 1 µm MyOne beads and a second independent magnetic tweezer setup (Supporting Information Section S7). These experiments demonstrate that the software performs robustly across different bead types and experimental configurations.

Once suitable beads have been selected, the final calibration curve can be generated via the “get master force calibration curve” panel and exported directly from the GUI. CALISTO generates a YAML file containing the fitted calibration parameters and a reference table of the force-magnet position relations, a Pickle archive containing the processed data and analysis results, and a CSV file for convenient use in external software environments (more details about the file formats are provided in the GitHub repository github.com/lipfertlab/calisto).

### Assessment of New Implementations

To evaluate the performance of CALISTO, we systematically investigated how several analysis and quality-control steps affect force calibration accuracy and robustness. Calibration quality was quantified using the root mean square error (RMSE), cross-validated RMSE, and the reduced χ^2^ statistic (Materials and Methods).

We first evaluated the effect of selecting only high-quality reference beads for drift correction. In this comparison, calibrations obtained using all manually identified reference beads were compared with calibrations obtained after applying CALISTO’s semi-automated reference-bead selection procedure. As described above, this procedure excludes reference beads that exhibit unusually large fluctuations, drift, sudden position jumps, or other statistical properties that differ substantially from the majority of reference traces. Restricting the analysis to these high-quality reference beads resulted in a modest but consistent improvement in calibration accuracy. The in-sample RMSE decreased by 3.7%, while the cross-validated RMSE and reduced χ^2^ remained essentially unchanged (Table 2). These results suggest that the selected reference beads improve data quality without introducing overfitting. We note, however, that the relatively small improvement observed here likely reflects the already high quality of the underlying dataset. In lower-quality datasets, unsuitable reference beads can substantially increase uncertainties and lead to noticeable changes in the fitted calibration parameters (Supporting Information Section S7).

**Table 2:** Effect of reference bead quality, motor-detection parameters, and plateau filtering on force– magnet position calibration performance. Calibration quality is quantified by the in-sample root-mean-square error (RMSE), cross-validated RMSE (CV RMSE), and reduced chi-squared 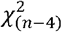 of the multiplicative bi-exponential fit. Applying the plateau filter substantially reduces residual scatter, while variations in bead quality and motor-detection parameters produce smaller differences in predictive performance.

| Reference bead | Tol./ plat. length | Plateau filter | RMSE [pN] | CV RMSE [pN] | $\chi^2_{n-4}$ |
| --- | --- | --- | --- | --- | --- |
| High-quality | High-quality | Not applied | 2.97 | $2.78 \pm 1.61$ | 1.002 |
| Low-quality | High-quality | Not applied | 3.08 | $2.78 \pm 1.26$ | 1.002 |
| High-quality | Low-quality | Not applied | 3.42 | $3.02 \pm 1.59$ | 1.002 |
| High-quality | High-quality | Applied | **1.91** | **$1.94 \pm 0.19$** | **1.003** |
| Low-quality | High-quality | Applied | 1.93 | $2.01 \pm 0.42$ | 1.004 |
| High-quality | Low-quality | Applied | 1.94 | $2.06 \pm 0.24$ | 1.005 |

Next, we evaluated the identification of stationary magnet positions. Force calibration relies on measurements acquired while the magnets are not moving and the bead-tether system has reached equilibrium. CALISTO therefore partitions the data into moving and stationary regions based on the motor position data. Adjusting the motor-movement detection parameters to more accurately identify stationary regions reduced calibration scatter, decreasing the in-sample RMSE by 12.9% and the cross-validated RMSE by 10.2% (Table 1). This result highlights the importance of correctly identifying force plateaus before any further analysis is performed.

The largest improvement was obtained from the automated detection of stable force plateaus. Here, the comparison is not based on motor position, but on the statistical properties of the fluctuations within each plateau. Even when the magnets are stationary, some plateaus contain slow drift, tracking artifacts, or other nonstationary behavior that violates the assumptions underlying variance-based force estimation. To identify such cases, CALISTO analyzes the dominant noise characteristics of each plateau using the lag-1 autocorrelation method and estimates the corresponding power-law spectral exponent. Only plateaus whose noise behavior falls within the range for which the uncertainty of the Allan variance or Hadamard variance is known analytically are retained for further analysis.

Applying this filtering step reduced the number of analyzed plateaus from n = 966 to n = 342. Despite this substantial reduction in dataset size, calibration performance improved remarkably. The filtered dataset exhibited a 36% reduction in RMSE and a 30% reduction in cross-validated prediction error (Table 2). Residuals remained centered around zero and showed no systematic dependence on magnet position, indicating that the procedure reduces noise without distorting the underlying force– distance relationship (Supporting Information Section S7).

To test whether removing these plateaus introduced bias, we compared calibration curves obtained from the full dataset and from the filtered dataset. The two independently fitted force–magnet position relations differed on average by only 0.145 pN across the entire calibration range. This difference is small compared to the overall force range and is fully consistent with statistical variability expected from random subsampling. Additional null-model analyses that account for unequal sample sizes and random label assignment yielded similar results. Together, these findings demonstrate that the filtering procedure improves precision while preserving the underlying force-magnet height relation.

Importantly, the same trends were observed for both MyOne and M270 beads and were reproduced on an independent magnetic tweezers instrument (Supporting Information Section S7), indicating that the benefits of the method are not specific to a particular bead type or experimental setup.

Overall, these results show that CALISTO’s quality-control procedures improve calibration accuracy at multiple stages of the workflow. While selecting suitable reference beads and accurately identifying stationary magnet positions provide measurable gains, the largest improvement arises from filtering out plateaus with unsuitable noise characteristics. This approach substantially reduces statistical uncertainty while preserving the underlying force–magnet position relationship, resulting in more robust and reproducible force calibration.

## CONCLUSION

In this study, we introduced and validated CALISTO, an open-source software framework for force calibration in multiplexed magnetic tweezers experiments. Designed to promote accessibility, reproducibility, and transparency, CALISTO combines automated analysis routines with user-guided quality control in a modular workflow. The software integrates key calibration steps, including reference bead selection, drift correction, offset determination, tether classification, force estimation, and generation of force–magnet position calibration curves. In addition to established calibration approaches based on the power spectral density (PSD) and Allan variance (AV), CALISTO implements our recently developed Hadamard variance (HV) method, which offers improved robustness against drift and correlated noise.

A central finding of this work is that automated identification of stable force plateaus substantially improves calibration precision. Unlike conventional approaches that analyze all stationary magnet positions equally, CALISTO additionally evaluates the statistical properties of the fluctuations within each plateau. Plateaus dominated by drift, tracking artifacts, or other nonstationary noise sources are automatically identified and excluded. This filtering reduces residual scatter, lowers prediction error, and improves the stability of the resulting calibration curves. Importantly, comparison of the calibration curves before and after filtering shows that the underlying force–magnet position relationship remains unchanged within statistical uncertainty, demonstrating that the procedure reduces noise without introducing measurable bias.

We further evaluated the influence of several user-adjustable analysis parameters. Improved identification of stationary magnet positions and the use of high-quality reference beads resulted in modest but measurable improvements in calibration accuracy. The largest gains, however, arose from the noise-based plateau filtering procedure. Together, these results highlight the importance of careful quality control throughout the calibration workflow.

Our analysis also demonstrates the importance of appropriate statistical fitting procedures. Because calibration residuals increase with force, weighted fitting approaches that account for the force dependence of the bead-to-bead variability provide more reliable parameter estimates across the full force range than unweighted methods. While both mono-exponential and bi-exponential functions can provide useful empirical descriptions of force–magnet position relationships, model selection should be guided by the data rather than by a priori assumptions. CALISTO therefore provides multiple fitting options together with quantitative metrics, such as the Akaike Information Criterion (AIC), to facilitate objective model comparison.

Overall, CALISTO provides a standardized and reproducible framework for magnetic tweezer force calibration. By combining automated quality control, transparent data processing, and quantitative performance evaluation, the software improves calibration robustness while remaining adaptable to different bead types, instrument designs, and experimental protocols. We anticipate that CALISTO will facilitate more consistent force calibration across laboratories and support the growing use of high-throughput single-molecule force spectroscopy.

## AUTHOR CONTRIBUTIONS

S.D.P. and A.U. share first authorship: they designed and executed the study, developed analytic tools, analyzed the data, and drafted the manuscript. C.K. conducted experiments, performed alpha testing of CALISTO, and contributed to writing and editing. L.R.S and K.R.S. conducted experiments. M.V. contributed to software testing, validation, and debugging. L.F. assisted with manuscript preparation and supervision, and secured funding. J.L. co-authored the manuscript and secured funding.

## ACKNOWLEDGMENTS

This research was financed by the European Research Council (Consolidator Grant “ProForce” 101002656), the Dutch Research Council (NWO, grant OCENW.GROOT.2019.071), the NWO-XL Open Competition Domain Science Grant (OCENW.XL.23.150), and the Horizon Europe 2021-2027 Framework Programme Grant (Agreement number 101168851). S.D.P. also received funding from the Alexander von Humboldt Foundation through a Feodor-Lynen Fellowship. The authors gratefully acknowledge Dave van den Heuvel, Elleke van Harten, Maria Kurilova, Sidonie Lieber, Andreas Spörhase, and Olga Ustinov for their valuable laboratory assistance, and Nynke Dekker, David Dulin, and John van Noort for fruitful discussions.

## DATA AVAILIBILITY

CALISTO is openly available and can be downloaded from github.com/lipfertlab/calisto. We also included information about file structure and data requirements as well as a manual and sample datasets for testing. The raw data analyzed in this study will be uploaded to zenodo upon publication.

## Declaration of generative AI and AI-assisted technologies in the manuscript preparation process

During the preparation of this work, the author(s) used GitHub copilot for coding assistance. The author(s) reviewed and edited the output as needed and take full responsibility for the content of the published article.

