## Supplementary Information for "CALISTO: A Standardized and Open-Source Software for Magnetic Tweezer Force Calibration"

### S1. Benchmarking of z-Offset Determination Procedures

To evaluate the accuracy of our z-offset detection procedure using an extension trace where the force is lowered to ~0 pN (Fig. S1A), we compared the Gaussian mixture model (GMM)-based method with several independent approaches. As a reference, we use the z-offsets determined by inducing a large number of plectonemic supercoils using magnet rotations, to decrease the tether extension to near zero (Fig. S1B).

In the first method, we applied a GMM to the vertical (z) position distribution of the tethered bead extension data, as described in the main text (Fig. S1A). As a physical reference point, we used magnet rotation (Fig. S1B): by incrementally rotating the magnets, we change the linking number of the DNA molecule. For torsionally constrained (i.e., supercoilable) tethers, this imposed twist accumulates as torsional stress, which is eventually relieved by the formation of plectonemes. The appearance of plectonemes leads to a progressive reduction in the bead's vertical position. As rotation continues, the tether shortens until the bead reaches a plateau corresponding to the maximally supercoiled state, where the bead contacts the surface. We define the rotation trace offset as the mean bead extension within this plateau region, i.e., the state of minimal extension.

We find that the offset values obtained from the GMM closely match the average extension measured at the surface-contact point during the rotation assay. This agreement demonstrates that the GMM-based approach reliably estimates the zero-offset point without requiring active rotation.

To evaluate method robustness across multiple tethers, we analyzed 17 supercoilable DNA molecules and calculated the mean and standard error of the estimated offsets using the GMM, minimum (after smoothing, see main text for details) and rotation-based methods. We also compared these with two alternative strategies: a piecewise-continuous trapezoidal (Trap) function fit and a fit to a Gumbel distribution.

The Trap function is designed to approximate the shape of the bead's z-position trace, consisting of a central plateau connected by linear transitions to zero (Figure 3B, main text). It is defined as:

$$f(x) = \begin{cases} L, \ |x-c| \le w_c/2 \\ \frac{L}{w_l}\left[x - \left(c - \frac{w_c}{2} - w_l\right)\right], \ -w_l \le x - c + \frac{w_c}{2} < 0 \\ \frac{L}{w_r}\left[x - \left(c - \frac{w_c}{2} - w_r\right)\right], \ 0 \le x - c - \frac{w_c}{2} < w_r \\ 0, \ \text{otherwise} \end{cases} \tag{1}$$

where $L$ is the plateau height, $c$ its center (used as the offset estimate), $w_c$ the width of the central region, and $w_l$, $w_r$ the widths of the left and right transition regions. We fit this function via least squares to the z-position distribution.

In parallel, we employed a Gumbel distribution fit, which is particularly suited to asymmetric distributions such as those observed at low forces where extension noise is skewed. The probability density function is:

$$P(x|\mu,\beta) = \frac{1}{\beta}\exp\left(-\frac{x-\mu}{\beta} - e^{-\frac{x-\mu}{\beta}}\right), \tag{2}$$

where $\mu$ is the location parameter (taken as the offset estimate) and $\beta$ is the scale. We applied maximum likelihood fitting to the low-extension region identified using the Trap method. Comparing all approaches, we find that the GMM, minimum and rotation-based methods yield consistent mean z-positions within experimental uncertainty (Fig. S1C). The Trap and Gumbel methods systematically underestimate the offset, likely due to their sensitivity to distribution tails and plateau width definitions.

Importantly, although the rotation-based approach provides a direct, physically grounded measurement of the offset corresponding to a well-defined mechanical transition, it is only applicable to supercoilable (i.e., torsionally constrained) DNA tethers. Tethers with single-stranded nicks or gaps cannot support torsional stress and thus do not form plectonemes under rotation. As a result, the rotation-based method cannot be used in such cases. In contrast, the GMM-based method does not rely on supercoiling and can be applied uniformly to both constrained and nicked tethers. Its combination of general applicability, model-free implementation, and agreement with rotation-based measurements makes it the most suitable routine for automated offset determination across diverse tether conditions.

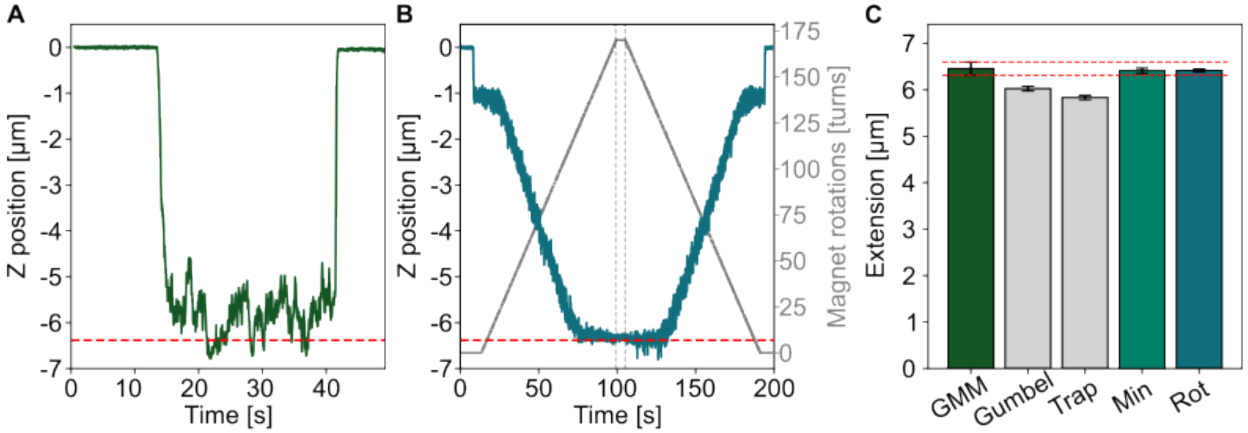

**Figure S1: Comparison of offset detection methods. (A**) Gaussian mixture model (GMM) fit to bead height distribution (green). The red dashed line represents the offset value determined from the GMM fit. **(B)** Bead extension (petrol) as a function of magnet rotation (grey). For torsionally constrained tethers, added turns induce plectoneme formation, leading to a progressive decrease in extension. The offset is defined as the rotation angle at which the bead reaches the surface. **(C)** Extension values determined across 17 supercoilable tethers using the GMM, rotation-based, minimum-extension, Trap (trapezoidal fit), and Gumbel fit methods. GMM, minimum and rotation offsets agree within experimental error, while Trap and Gumbel methods underestimate, the offset. The red dashed lines represent the standard errors of the GMM mean.

### S2. Bead-to-Bead Variations of M270 Magnetic Beads

To quantify the relative bead-to-bead deviation from the mean force $F_{mean}$ we calculate the bead-to-bead force deviations from the force data presented in the main text (Fig. 4) and fit the data (Fig. S2) with $\sigma_{Fmean} = \sigma_0 + \sigma_{rel}F$, where $\sigma_{Fmean}$ represents the bead-to-bead deviation from the mean force in pN, $F_{mean}$ (in pN) the mean force, and $\sigma_{rel}$ the unitless, relative bead-to-bead deviation from the mean force. The error $\sigma_0$ (in pN) accounts for intrinsic system errors. The derived values for $\sigma_{rel.}$ and $\sigma_0$ are 8.6 % and 0.07 pN, respectively. The relative error of ~9% is fully consistent with previously observed bead-to-bead variations for these types of beads[1, 2], which dominate the force uncertainty for accurate measurements. The parameter $\sigma_0$ essentially estimates the "noise floor", i.e. the level of absolute uncertainty unrelated to bead-to-bead variability.

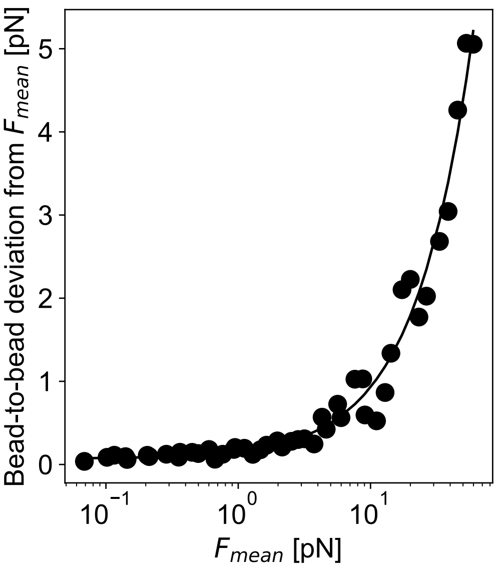

**Figure S2:** Experimental bead-to-bead deviations from the mean force as a function of the mean force derived from experimental data traces using 21 kbp dsDNA and M270 beads.

**S3. Tracking Frequency and Measurement Time Guideline for WLC Tethers**

To estimate the minimum required tracking frame rate and, in addition, the time required to sufficiently sample the bead fluctuations for a tethered bead system, we provide a python script (see gitub repository: github.com/lipfertlab/calisto) that calculates the characteristic or corner frequency of a given bead–tether system. To avoid undersampling, the acquisition frequency should be at least twice the characteristic frequency of the bead-tether system, which is known as the Nyquist frequency. If the sampling rate falls below this value, bead fluctuations are undersampled and high-frequency information is irreversibly lost. The Nyquist frequency therefore defines the lower bound for selecting a frame rate that fully resolves the mechanical response of the tether–bead system.

Here, we assume the tether to have a WLC response, which is a good approximation for both double-stranded DNA and unfolded proteins (albeit with different parameters). The force–extension behavior of the tether is described using the seven-parameter interpolation formula of the worm-like chain (WLC) model.[3] For a molecular extension $z$, persistence length $L_p$, contour length $L_c$, and absolute temperature $T$, the force is given by

$$F(z) = \frac{k_B T}{L_p}\left[\frac{1}{4(1-z/L_c)^2} + \sum_{p=0}^{7} a_p \,(z/L_c)^p\right], \tag{3}$$

where $k_B T$ is expressed in pN·nm and $a_p$ are the coefficients reported by Bouchiat *et* al[3] (see also main text).

The corresponding extension at a given applied force is then obtained by numerically solving $F(z) - F_{\text{applied}} = 0$ within the interval $0 < z < L_c$, ensuring stability near full extension. The effective longitudinal stiffness is defined as $\kappa(F) = F/z(F)$. In the high-force regime, where the extension approaches the contour length ($z \to L_c$), this expression admits the asymptotic approximation $\kappa(F) \approx F/L_c$. Assuming overdamped dynamics, the drag coefficient of a spherical bead with radius $R$ in a fluid of viscosity $\eta$ is calculated using Stokes' law, $\gamma = 6\pi\eta R$. The corner frequency follows from $f_c = \kappa(F)/(2\pi\gamma)$, and the Nyquist frequency is defined as $f_{\text{Nyquist}} = 2f_c$.

Figure S3.1 illustrates the minimum sampling frequency as a function of the force for the two magnetic bead types used in this study, i.e. MyOne (1 µm) and M270 (2.8 µm). The WLC tethers are 21kbp dsDNA strands with a contour length $L_c$ = 7140 nm and a persistence length $L_p$ = 45 nm. The temperature is 293 K. Note that this procedure is general and can be applied to any system that is well described by the worm-like chain (WLC) model.

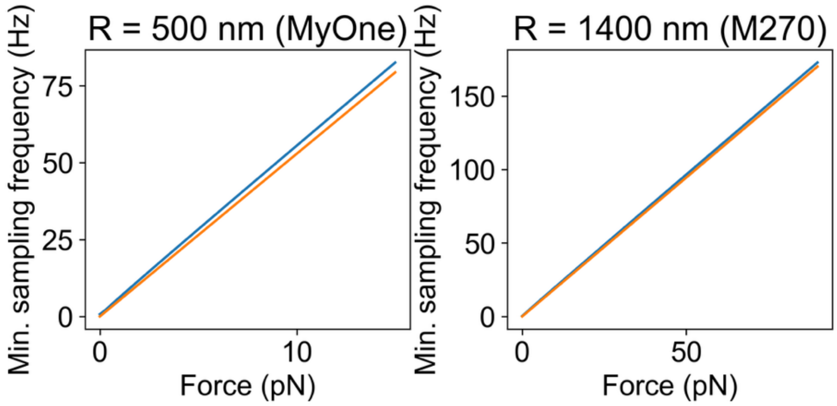

**Figure S3.1: Minimum sampling frequency as function of force.** The system consists of MyOne (left) and M270 (right) magnetic beads tethered by a 21 kbp double-stranded DNA molecule with a contour length $L_c = 7140$ nm and a persistence length $L_p = 45$ nm. The temperature is 293 K. Solid blue lines: WLC model; orange lines: asymptotic approximation (2 $f_c$).

The minimum measurement time required for reliable sampling can also be estimated from the corner frequency derived from the worm-like chain (WLC) stiffness and overdamped dynamics of the bead. The characteristic relaxation time of the system is given by $\tau_\mathrm{c} = (2\pi f_c)^{-1}$, where $f_c$ is the corner frequency. The number of statistically independent measurements is then approximately $N_\mathrm{ind} = T_\mathrm{meas}/\tau_\mathrm{c}$, with $T_\mathrm{meas}$ being the total measurement duration. Equivalently, the minimum measurement time required to obtain a target number of independent measurements can be estimated as $t_\mathrm{min} = N_\mathrm{ind}/(2\pi f_c)$.

The number of independent measurements determines the expected statistical uncertainty of the sampled trace. Following the argument by Strick et al.[4], the relative statistical error scales as $\varepsilon_\mathrm{rel} \propto \sqrt{\tau_\mathrm{c}/T_\mathrm{meas}}$. Figure S3.2 shows the recommended measurement times using $N_\mathrm{ind} = 1000$.

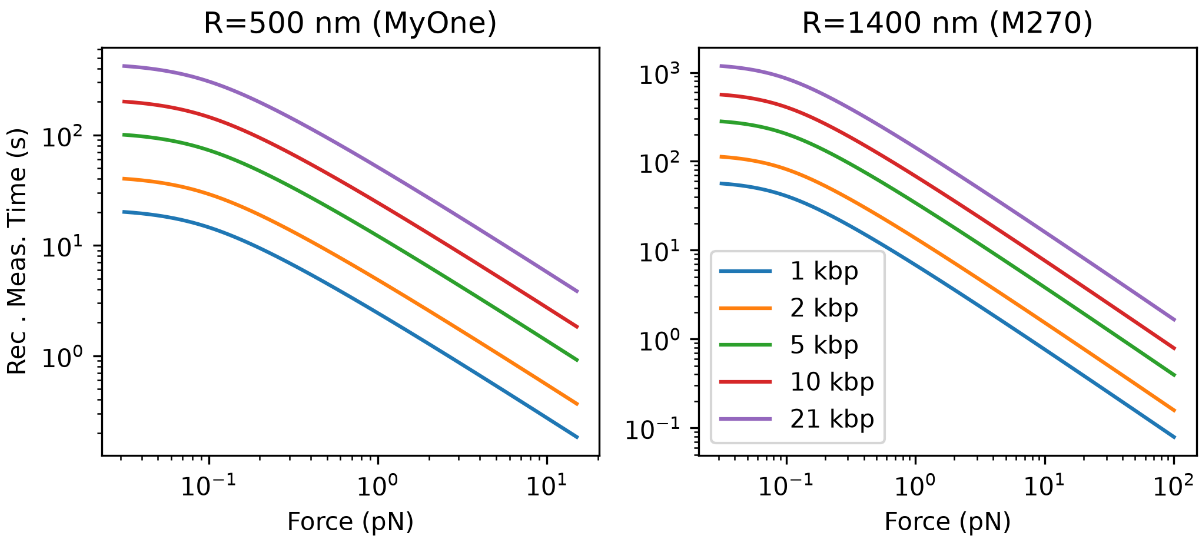

**Figure S3.2: Recommended measurement times as function of force.** The system models MyOne (left) and M270 (right) magnetic beads tethered by a double-stranded DNA molecule with various contour lengths (shown in the legend) and a persistence length $L_p = 45$ nm. The temperature is 293 K.

### S4. Tracking Resolution of Magnetic Tweezer

To quantify the tracking accuracy, we monitored the x, y, and z positions of surface-immobilized 1 µm polystyrene reference beads at frame rates of 58 Hz and 168 Hz. The positional noise (or tracking error) was determined from the standard deviation of the bead motion obtained via histogram analysis. To remove slow baseline drift, the time traces were detrended using a second-order Savitzky–Golay filter with a window length set to eight times the lag time corresponding to the minimum of the Allan deviation, i.e. 70 s (Fig. S4 A,B). The histogram of the detrended position data was fitted with a Gaussian function, yielding a standard deviation of σ ≈ 1 nm. Figure S4C summarizes the average standard deviations obtained from 36 beads at 58 Hz and 19 beads at 168 Hz, shown separately for the horizontal position (mean of x and y) and the vertical (z) position. Depending on tracking frequency and spatial direction, the standard deviation ranges from approximately 0.5 nm to 4.0 nm.

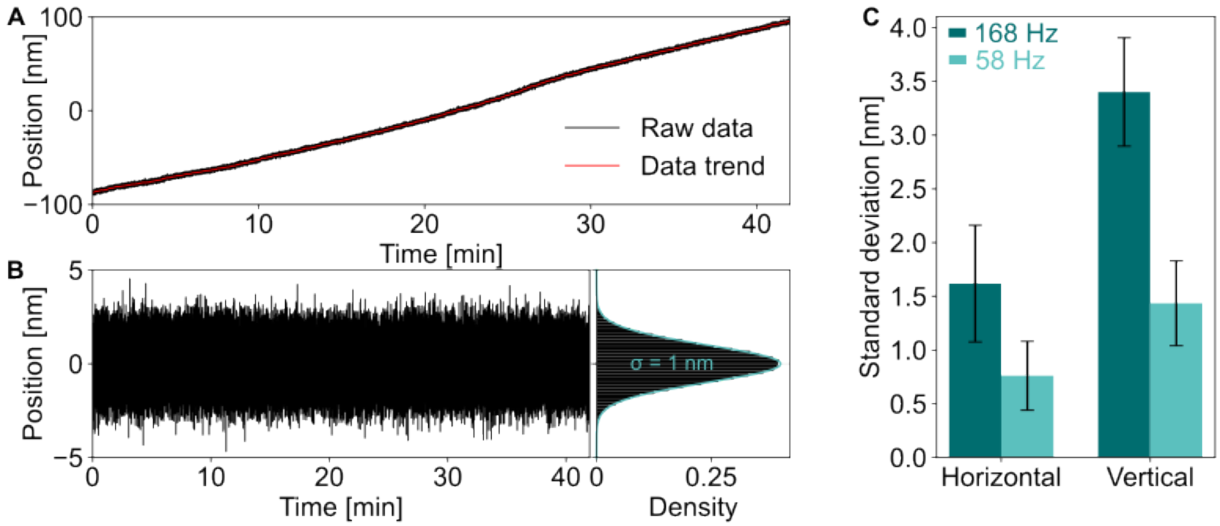

**Figure S4. Determination of tracking precision using a surface-immobilized reference bead.** (A) Raw time trace of the z position of a surface-immobilized 1 µm-diameter polystyrene bead (black) recorded at 58 Hz. The red curve shows the low-frequency trend of the signal. (B) Left: Detrended z-position trace obtained using a second-order Savitzky–Golay filter to remove baseline drift. Right: Histogram of the detrended data. The distribution is fitted with a Gaussian distribution (petrol), yielding a standard deviation of σ = 1 nm. (C) Mean standard deviations obtained from multiple reference beads tracked at 58 Hz (N = 36, petrol) and 168 Hz (N = 19, teal). The horizontal precision corresponds to the mean of the x and y positions, and the vertical precision to the z position. Tracking at the two different camera frequencies was carried out under otherwise identical illumination conditions. Consequently, for the higher frequency data fewer photons are recorded per frame, giving rise to the larger frame to frame standard deviation. We note that the ratios of the observed tracking accuracies are close to the expected value of $(168\text{ Hz}/58\text{ Hz})^{1/2}$ ~ 1.7.

**S5: Relation Between Degree of Off-Center Attachment and Force Uncertainties**

In magnetic tweezers (MT), the tether extension is determined from the z-position of the magnetic bead. If the tether is attached off-center to the bead, the measured tether extension is underestimated. Consequently, in a force calibration experiment, the force is also underestimated.

Suppose as shown in Fig. S5.1 that a bead of radius $R_B$ traces a circle with radius $R$ during a rotation test (see Fig. 3D of the main text). The resulting underestimation of the tether extension, $\Delta z$, can be obtained from the Pythagorean theorem,

$$\Delta z = R_B\left(1 - \sqrt{1 - (R/R_B)^2}\right). \tag{4}$$

Since the force is estimated as $F = \kappa L_{ext}$, the relative force error caused by the underestimated tether extension is

$$\varepsilon_{rel} = 1 - \frac{\kappa(L_{ext} - \Delta z)}{\kappa L_{ext}} = \frac{\Delta z}{L_{ext}} = \frac{R_B}{L_{ext}}\left(1 - \sqrt{1 - (R/R_B)^2}\right). \tag{5}$$

For example, for an M270 bead and a tether extension of 6 $\mu\text{m}$, corresponding to approximately $1\ \text{pN}$, a rotation radius of $R = 530\ \text{nm}$ leads to a relative force error of about 5%. A broader range of relative force errors can be found in Fig. S5.2.

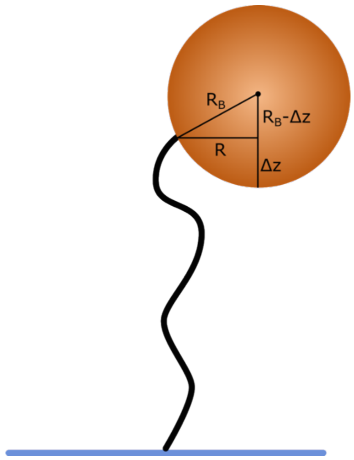

**Figure S5.1. Schematic of a magnetic bead with an off-center tether attachment.** If the tether is attached at a lateral distance $R$ from the bead center, the measured tether extension is underestimated by $\Delta z$. For a bead with radius $R_B$, this underestimation follows from the right triangle formed by $R$, $R_B - \Delta z$, and $R_B$.

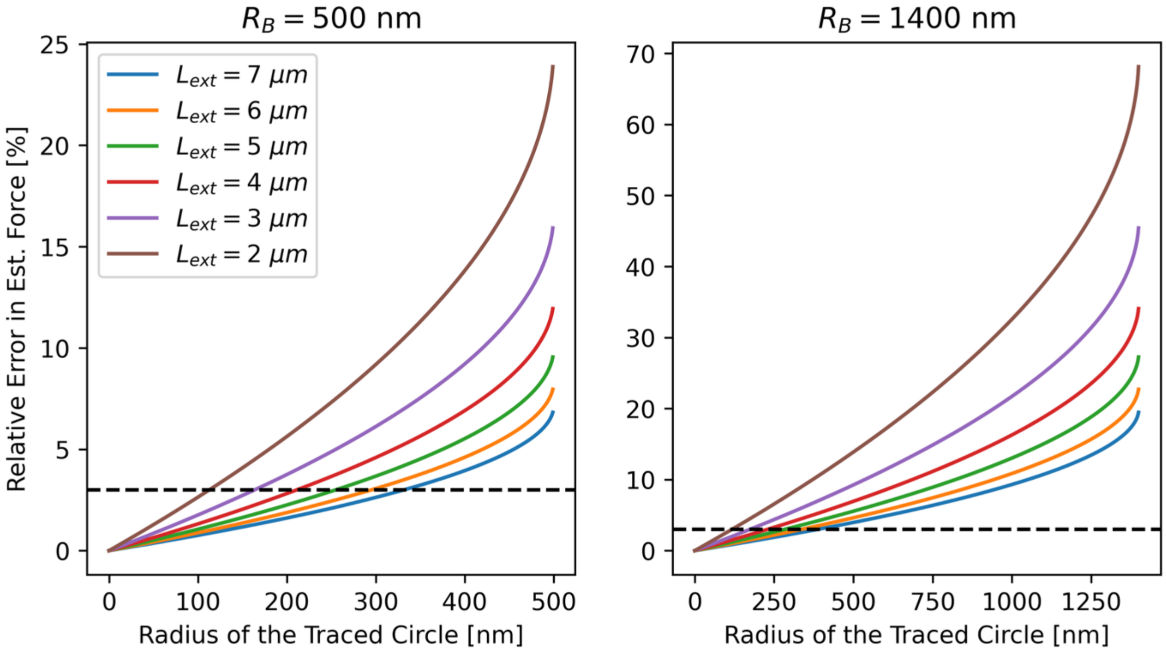

**Figure S5.2.** Relation between the radius of the circle traced by a magnetic bead during a rotation test and the relative error in the estimated force caused by off-center tether attachment. The left and right panels show the relation for bead radii of 500 nm and 1400 nm, corresponding to MyOne and M270 beads, respectively, for different tether extensions, $L_{\text{ext}}$. The black dashed line marks a relative error of 3%.

### S6. Force Calibration and Statistical Analysis of M270 Beads Using two MT Devices

We compare four fitting functions using force calibration data recorded from tethers composed of 21 kbp DNA attached to M270 magnetic beads. The functions are a mono-exponential ($F(z) = Ae^{-z/\lambda}$), a bi-exponential ($F(z) = F_{max}(ce^{-\frac{z}{\lambda_1}} + (1-c)e^{-\frac{z}{\lambda_2}})$), and multiplicative mono- and bi-exponential function presented in the main text. Here, $z$ denotes the magnet position measured relative to the sample plane, with $z = 0$ defined as the contact point between the lower surface of the magnets and the coverslip of the sample. In our setup, this corresponds to approximately 270 μm from the bottom slide of the flow cell. In total, we analyze 26 individual beads and evaluate the performance of each model using standard statistical metrics (Fig. S6). Model quality is evaluated from log-space residuals $r = \log(F) - \log(\hat{F})$ under a Gaussian likelihood. The mono-exponential model yields an Akaike information criterion (AIC) of 1249.49, and a Bayesian information criterion (BIC) of 1259.27. The bi-exponential shows an AIC = 1243.20 and a BIC = 1262.78. The multiplicative mono- and bi-exponential functions lower the AIC and BIC to $AIC_{mono}$ = -862.06 and $AIC_{bi}$= -1158.10 and $BIC_{mono}$ = -852.27.06 and $BIC_{bi}$= -1138.53, respectively. Since the simple and multiplicative exponential models have the same parameter count, the substantially lower information criteria of the multiplicative bi-exponential function arise from a markedly improved likelihood, indicating a superior description of the data in log space.

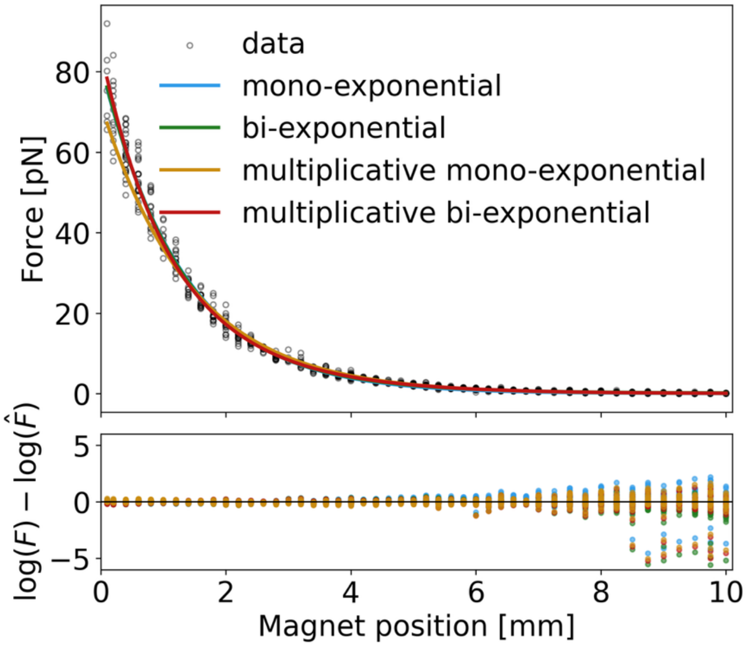

**Figure S6: Comparison of exponential models for force–magnet-position calibration.** Top: Measured force as a function of magnet position for a representative 21 kbp DNA tether attached to an M270 bead (open black circles), together with four fit functions, i.e mono-exponential (blue), bi-exponential (green), multiplicative mono-exponential (orange), and multiplicative bi-exponential (red.) Bottom: Log-space residuals, $\log(F) - \log(\hat{F})$, for the four models plotted against magnet position.

### S7. Force Calibration and Statistical Analysis of M270 and MyOne Beads Using two MT Devices

*Calibration quality:* Calibration quality was quantified using RMSE, cross-validated RMSE, and reduced $\chi^2$ statistics (see main text, Materials and Methods). We first assess the impact of restricting the calibration to high-quality reference beads. The fitting parameters derived for the M270 data presented in the main text is summarized in Table S1. The modest reduction of the in-sample RMSE by 3.4% can be explained by the lack of a very low-quality reference bead. If we purposefully include such a very low-quality bead, the RMSE reduced by 5% and the fitting parameters changed drastically ($\Delta F$=20%, mono -> bi-exponential).

**Table S1:** Summary of fitting parameters using the multiplicative double exponential fit described in the main text (Eq. (9)) and 21 kbp DNA. *Note: plateau filtering has not been applied*

| sample | Reference bead | $F_{max}$/ pN | c | $\lambda_1/\mu$m | $\lambda_2/\ \mu$m | RMSE | CV RMSE | $\chi^2$ |
|---|---|---|---|---|---|---|---|---|
| M270 | High-quality | 81.9 | 0.57 | 1.03 | 1.68 | 2.97 | 2.78 | 1.002 |
| M270 | Mid-quality | 79.1 | 0.57 | 1.07 | 1.69 | 3.08 | 2.78 | 1.002 |
| MyOne | High-quality | 12.99 | 0.81 | 1.01 | 2.35 | 0.89 | 0.92 | 1.004 |
| MyOne | Mid-quality | 13.62 | 0.72 | 0.94 | 2.11 | 0.72 | 0.80 | 1.002 |

*Statistical Tests:* To determine whether data filtering introduces bias or artifacts that alter the underlying force–magnet position relationship, rather than simply reducing force uncertainties, we performed two complementary statistical tests, i.e. Size-matched subsampling test and Permutation test (see main text, Materials and Methods for details).

To quantify whether the reduced dataset alters the inferred calibration curve, we first compute the normalized absolute curve difference *D* between the two fitted relationships given as

$$D = \frac{1}{x_{\max}-x_{\min}} \int_{x_{\min}}^{x_{\max}} \mid F_B(x) - F_A(x) \mid \, dx, \tag{6}$$

which represents the mean absolute force difference between the two calibrations across the shared magnet-position range and therefore has units of force (Fig S7.1B). In our data, the observed separation is $D_{\text{obs}} = 0.145$ pN, indicating that, on average, the inferred force responses differ by only a fraction of a piconewton across the entire domain.

To determine whether this difference exceeds what is expected from unequal sample size alone, we benchmark $D_{\text{obs}}$ against the two null models:

First, the larger dataset ($n_A = 1861$) is repeatedly subsampled to match the size of the filtered dataset ($n_B = 985$), yielding a size-matched null distribution $D_{\text{sub}}$ that captures variability arising purely from sampling. The probability $P(D_{\text{sub}} < D_{\text{obs}}) = 0.032$ shows that observed difference is significantly smaller than the difference caused by random dataset size reduction and therefore do not indicate systematic bias.

Second, the permutation test is performed by randomly reassigning dataset labels while preserving sample sizes, generating a permutation null distribution $D_{\mathrm{perm}}$ under the hypothesis that both datasets originate from the same underlying relationship. The resulting permutation probability ($p_{\mathrm{perm}} = 0.999$) likewise indicates that the observed curve separation is statistically indistinguishable from random variability.

To further evaluate CALISTO, we performed additional measurements using MyOne magnetic beads (1 µm diameter) tethered to the 21 kbp dsDNA construct described in the main text. Experiments were conducted both on the magnetic tweezers (MT) setup used in the main text and on a second, independent MT instrument. The latter system uses a 650 nm LED (Thorlabs, Germany) for illumination and an Optronis CP80-25-M-72 camera (Optronis, Germany) operating at 72 Hz for bead tracking. Fitting the data with the multiplicative bi-exponential model described in the main text yielded maximum forces of 10.41 pN and 12.55 pN for the two setups, respectively (Fig. S7.2 A,C). In both cases, statistical analysis (size-matched null distribution and permutation test; see Materials and Methods) shows that filtering for stable plateaus does not introduce bias and does not alter the underlying force relationship (Fig. S7.2 B,C).

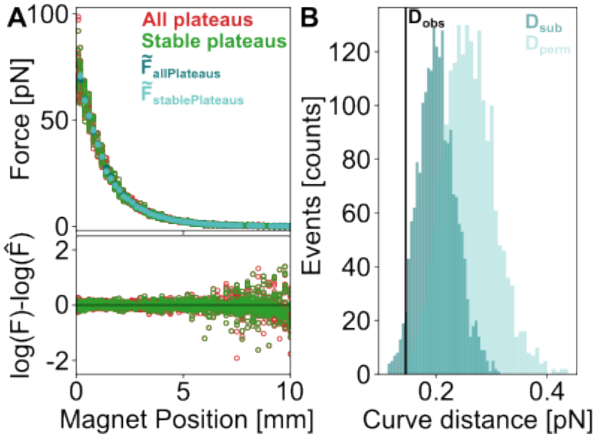

**Figure S7.1: Assessment of stable force plateau filtering and its impact on force calibration using M270 beads.** (A) Force–magnet position data obtained from plateau segments of bead position time traces of 30 independent beads. The full dataset ("All plateaus", red, $n = 1861$) is compared with the filtered dataset ("Stable plateaus", green, $n = 985$), where filtering is based on the lag-1 autocorrelation of bead-position fluctuations within each plateau. Solid lines show independent double-exponential fits to each dataset. $\tilde{F}_{allPlateaus}$ and $\tilde{F}_{stablePlateaus}$ denote binned forces defined as the median force within magnet-position bins and used to determine the absolute force difference (Eq. 6). The lower panel shows logarithmic residuals, $\log\, F(x) - \log\, \hat{F}(x)$, highlighting reduced scatter for the filtered data. (B) $D_{\mathrm{sub}}$(dark teal) is obtained from size-matched subsampling of the larger dataset and represents the variation expected from differences in dataset size alone. $D_{\mathrm{perm}}$(light teal) is obtained by random permutation of dataset labels and represents the variation expected if both datasets were drawn from the same underlying force–position relationship. The vertical solid line marks the observed value $D_{obs}$ of the absolute force difference between the two binned curves shown in panel A. Its position within both distributions indicates that the difference between filtered and unfiltered calibration curves is consistent with sampling variability rather than a systematic change in the underlying relationship.

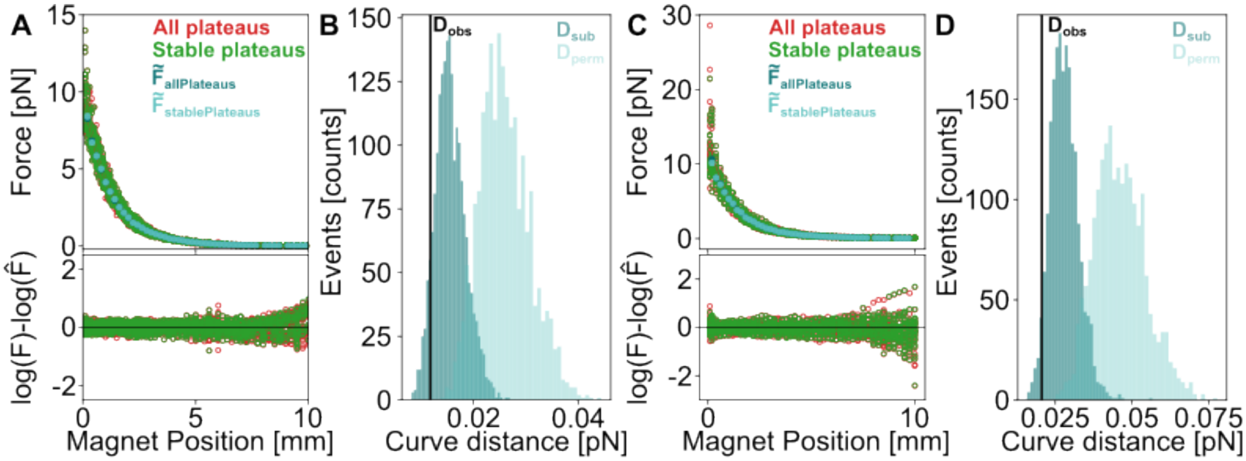

**Figure S7.2: Force calibration curves of MyOne magnetic beads obtained using 21 kbp DNA. (A,B)** Measurements performed with the MT device equipped with a Falcon camera at a sampling rate of 58 Hz. The maximum force reaches 10.31 pN when all plateaus are included and 10.41 pN when only stable plateaus are considered (A, top). The log-residuals are centered around zero (A, bottom). The observed curve distance is $D_{\text{obs}} = 0.012\text{pN}$, with $P(\text{distance} \geq \text{observed} \mid \text{size effect only}) \approx 0.9135$, and a permutation p-value $P(\text{distance} \geq \text{observed}) = 1$. **(C,D)** Measurements performed with the MT device using an Optronis camera at a sampling rate of 72 Hz. The maximum force is 12.99 pN when all plateaus are included and 12.55 pN when only stable plateaus are considered (C, top). The log-residuals remain centered around zero (C, bottom). The observed curve distance is $D_{\text{obs}} = 0.021\text{pN}$, with $P(\text{distance} \geq \text{observed} \mid \text{size effect only}) \approx 0.962$, and a permutation p-value $P(\text{distance} \geq \text{observed}) = 1$.
